# The active zone protein Clarinet regulates ATG-9 trafficking at synapses and presynaptic autophagy

**DOI:** 10.1101/2021.08.19.457026

**Authors:** Zhao Xuan, Sisi Yang, Sarah E. Hill, Benjamin Clark, Laura Manning, Daniel A. Colón-Ramos

## Abstract

In neurons, autophagy is temporally and spatially regulated to occur near presynaptic sites. How trafficking of autophagy proteins is regulated to support synaptic autophagy is not well understood. From forward genetic screens, we identify a role for the long isoform of the active zone protein Clarinet (CLA-1L) in regulating trafficking of autophagy protein ATG-9 at synapses, and presynaptic autophagy. ATG-9 is a transmembrane protein that undergoes activity-dependent exo-endocytosis at synapses, and mutations in CLA-1L result in abnormal accumulation of ATG-9 into clathrin-rich endocytic intermediates. CLA-1L extends from the active zone to the periactive zone, and genetically interacts with periactive zone proteins required for clathrin-dependent endocytosis. We find that CLA-1L is specifically required for sorting of ATG-9 at synapses, likely via endosome-mediated endocytosis, and for activity-dependent presynaptic autophagy. Our findings provide mechanistic insights into how active zone proteins regulate key steps of ATG-9 exo-endocytosis, a process that could couple the activity state of the neuron and autophagy.

**Highlights:**

• The long isoform of the active zone protein Clarinet (CLA-1L) regulates ATG-9 trafficking at synapses
• CLA-1L extends from the active zone to the periactive zone and cooperates with the periactive zone endocytic proteins EHS-1/EPS15 and ITSN-1/ intersectin 1 in ATG-9 trafficking during exo-endocytosis
• Mutations in CLA-1L, or in clathrin-associated adaptor molecules, result in abnormal accumulation of ATG-9 into clathrin-rich endocytic intermediates
• CLA-1L mutants which affect ATG-9 trafficking are also defective in activity-dependent presynaptic autophagy

## Introduction

Macroautophagy (herein called autophagy) is a well conserved cellular degradative pathway, and its disruption in neurons result in axonal degeneration, accumulation of protein aggregates and cell death^1–4^. Neuronal autophagy is regulated to cater to the particular physiological needs of neurons^5, 6^. Regulated steps in neurons include temporal and spatial control of autophagosome biogenesis near synaptic sites and in response to increased neuronal activity states^7–18^. The spatial regulation of autophagosome biogenesis at synapses results in part from directed transport of autophagic proteins to presynaptic sites and their local trafficking at the synapse^7, 11, 12, 19–21^.

Trafficking of autophagic proteins at synapses is mechanistically coupled to resident synaptic proteins^6, 22–24^. In both vertebrate and invertebrates, synaptic proteins endophilin and synaptojanin 1, best known for their roles in regulating the synaptic vesicle cycle, also play critical roles in the trafficking of autophagy proteins and the induction of synaptic autophagy^10, 21, 25, 26^. Bassoon, a large active zone protein associated with synaptic vesicle retention, was similarly implicated in the regulation of autophagy in neurons by interacting with the autophagy protein ATG-5^27^, and the E3 ubiquitin ligase Parkin^28, 29^. These mechanistic findings support a model by which the synaptic machinery, including structural active zone proteins, coordinate local synaptic autophagosome biogenesis through the local regulation of proteins involved in autophagy. Yet, how structural components of the synapses, such as active zone proteins, regulate autophagy at synapses is not well understood.

ATG-9 is the only transmembrane protein in the core autophagy pathway^30–34^, and is actively trafficked to promote local autophagosome biogenesis^35–39^, including that at synapses^12^. ATG-9 is transported to synapses in Golgi-derived vesicles, and ATG-9-containing vesicles undergo activity-dependent exo-endocytosis at synapses^21^. Disruption of ATG-9 trafficking at synapses, including its capacity to undergo exo-endocytosis, suppresses activity-dependent increases in autophagy at presynaptic sites^12, 21^. Therefore, ATG-9 trafficking at synapses is important for local autophagosome biogenesis.

To better understand the mechanisms that regulate ATG-9 trafficking at synapses, and local autophagy, we performed unbiased forward genetic screens *in vivo* in *Caenorhabditis elegans* (*C. elegans*) neurons. We find that the long isoform of the active zone protein Clarinet (CLA-1L), which bears functional similarity to Piccolo and Bassoon^40–42^, regulates ATG-9 trafficking at synapses, and presynaptic autophagy. In *cla-1(L)* mutants, ATG-9 abnormally accumulates into clathrin-rich presynaptic structures, which can be suppressed by mutants for synaptic vesicle exocytosis, suggesting that the ATG-9 phenotype in *cla-1(L)* mutants emerges from defects in ATG-9 sorting during exo-endocytosis. We observe that CLA-1L extends from the exocytic active zone to the endocytic periactive zone, and genetically interacts with the periactive zone proteins EHS-1/EPS15 and ITSN-1/intersectin 1 in mediating ATG-9 sorting at presynaptic sites. We also find that mutants of the clathrin-associated adaptor complexes AP-2 and AP180 phenocopy and enhance the ATG-9 phenotypes observed for *cla-1(L)* mutant, whereas mutants for the AP-1 adaptor complex and the F-BAR protein syndapin 1 suppress the phenotype. Disruptions of CLA-1L that result in abnormal ATG-9 localization also affect activity-dependent presynaptic autophagy.

Our findings support a model whereby CLA-1L bridges the exocytic active zone regions with the endocytic periactive zone to regulate presynaptic sorting of ATG-9, likely via endosome-mediated endocytosis. Our findings also underscore the importance of active zone proteins in regulating local trafficking of autophagy proteins and presynaptic autophagy, which could enable coupling of neuronal activity states with biogenesis of local synaptic autophagosomes.

## Results

### The active zone protein Clarinet (CLA-1) regulates ATG-9 trafficking at presynaptic sites

We examined the *in vivo* localization of ATG-9 in the AIY interneurons of *C. elegans*. AIYs are a pair of bilaterally symmetric interneurons which display a stereotyped distribution of presynaptic specializations along their neurites (Figure 1A and C)^43, 44^. Simultaneous visualization of ATG-9::GFP and the presynaptic marker mCherry::RAB-3 indicated that ATG-9 is enriched at presynaptic sites in AIY, consistent with previous studies (Figure 1 B-D)^12, 21^.

**Figure 1.**
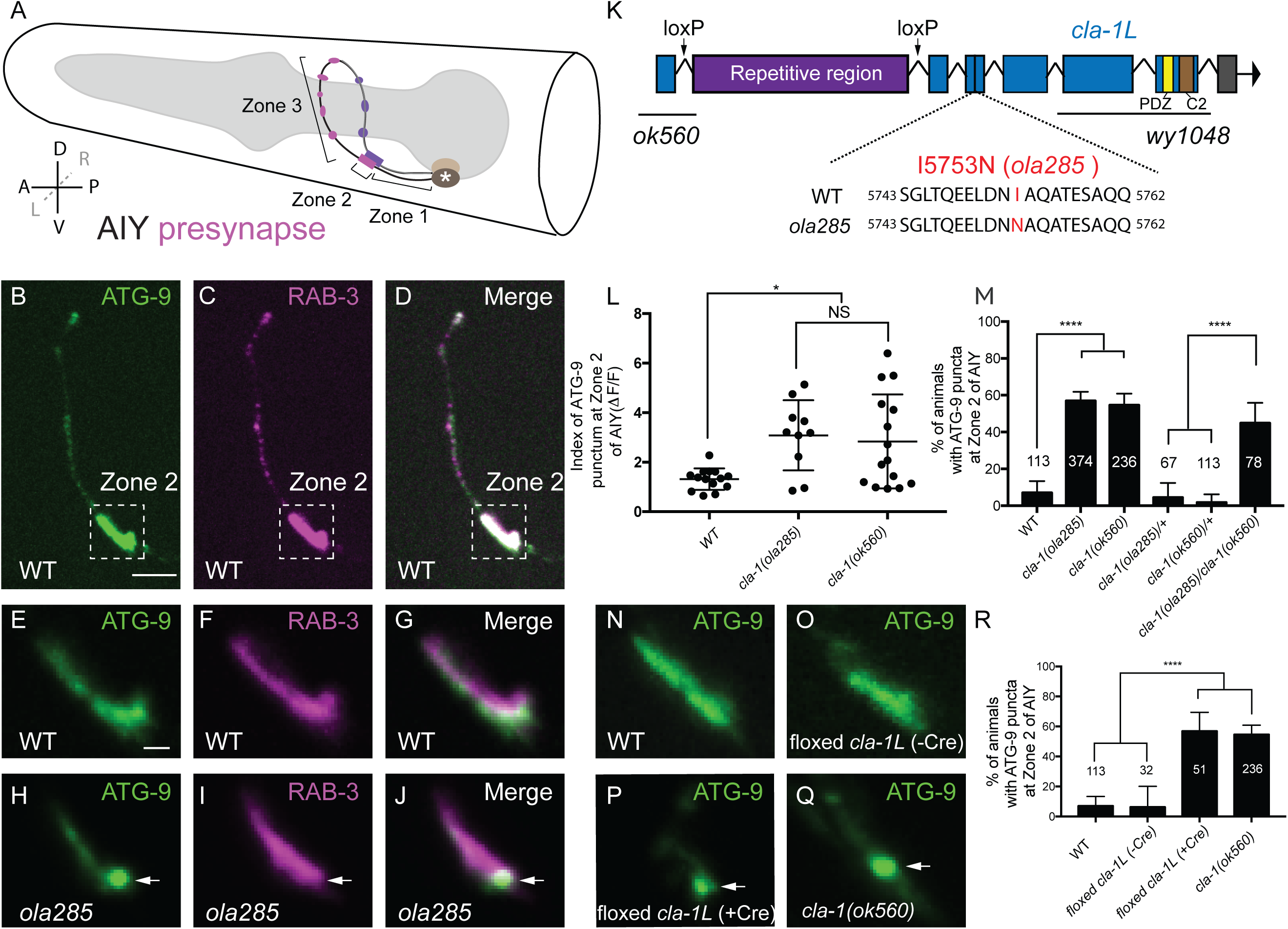
The long isoform of Clarinet (CLA-1L) regulates ATG-9 trafficking at presynaptic sites. (A) Schematic of the head of *C. elegans*, including pharynx (grey region) and the two bilaterally symmetric AIY interneurons. The asterisk denotes the cell body. There are three distinct segments along the AIY neurite: an asynaptic region proximal to AIY cell body (Zone 1), a large presynaptic region (Zone 2), and a segment with discrete presynaptic clusters at the distal part of the neurite (Zone 3)^43, 44^. Presynaptic regions (Zone 2 and Zone 3) are in magenta (AIYL) or violet (AIYR). In axis, A, anterior; P, posterior; L, left; R, right; D, dorsal; V, ventral. (B-D) Distribution of ATG-9::GFP (B) and synaptic vesicle protein (mCherry::RAB-3, pseudo-colored magenta) (C) in the synaptic regions of AIY (merge in D). The dashed box encloses AIY Zone 2. (E-J) Distribution of ATG-9::GFP (E and H) and synaptic vesicle protein (mCherry::RAB-3, pseudo-colored magenta) (F and I) at Zone 2 of AIY (merge in G and J) in wild type (WT) (E-G) and *ola285* mutant (H-J) animals. ATG-9 is evenly distributed in WT, but forms subsynaptic foci in *ola285* mutants, which are not enriched with RAB-3 (indicated by arrows in H-J). (K) Schematic of the genomic region of *cla-1L*. The locations of loxP sites and the genetic lesions of the *cla-1* alleles examined in this study are indicated. The genetic lesion in allele *ola285* (I to N at residue 5753) is shown for both WT and *ola285* mutants. The positions of the repetitive region in CLA-1L, and the conserved PDZ and C2 domains in all CLA-1 isoforms are also shown in the schematic. (L) Quantification of the index of ATG-9 punctum (ΔF/F; See Methods) at Zone 2 of AIY in wild type (WT), *cla-1(ola285)* and *cla-1(ok560)* mutants. Error bars show standard deviation (SD). “NS” (not significant), *p<0.05 by ordinary one-way ANOVA with Tukey’s multiple comparisons test. Each dot in the scatter plot represents a single animal. (M) Quantification of the percentage of animals displaying ATG-9 subsynaptic foci at AIY Zone 2 in the indicated genotypes. Error bars represent 95% confidence interval. ****p<0.0001 by two-tailed Fisher’s exact test. The number on the bars indicates the number of animals scored. (N-Q) Distribution of ATG-9::GFP at Zone 2 of AIY in wild type (WT) (N), *floxed cla-1L without Cre* (O) and *floxed cla-1L with Cre* expressed cell-specifically in AIY (P) and *cla-1(ok560)* (Q) animals. Arrows (in P and Q) indicate abnormal ATG-9 foci. (R) Quantification of the percentage of animals displaying ATG-9 subsynaptic foci at AIY Zone 2 in the indicated genotypes. Error bars represent 95% confidence interval. ****p<0.0001 by two-tailed Fisher’s exact test. The number on the bar indicates the number of animals scored. Scale bar (in B for B-D), 5 μm; (in E for E-J and N-Q), 1 μm.

To understand ATG-9 trafficking at the presynapse and its role in regulating presynaptic autophagy, we performed forward genetic screens for genes regulating presynaptic localization of ATG-9. From this screen, we identified an allele, *ola285*, which displays abnormal ATG-9 distribution at synapses. In wild-type animals, ATG-9 is evenly distributed in the presynaptic-rich region (termed Zone 2; Figure 1E-G and M), whereas in the *ola285* mutants, ATG-9 is enriched in subsynaptic foci in about sixty percent of animals (Figure 1H-J and M). To quantify the expressivity of the phenotype, we calculated an index (*Fpeak*-*Ftrough*)/*Ftrough* for ATG-9::GFP. The index was calculated from representative micrographs of the ATG-9 phenotype in the indicated genotypes (see Methods). Our quantifications of expressivity revealed a significant redistribution of ATG-9 to subsynaptic foci in *ola285* mutants as compared to wild-type animals (Figure 1L).

To identify the genetic lesion corresponding to the *ola285* allele, we performed single-nucleotide polymorphism (SNP) mapping coupled with whole genome sequencing^45–47^. We identified the genetic lesion of *ola285* in the locus of the gene *cla-1*, which encodes for Clarinet. Clarinet (CLA-1) is an active zone protein that contains PDZ and C2 domains with similarity to vertebrate active zone proteins Piccolo and RIM (Figure 1K)^40–42^. Three lines of evidence support that *ola285* is an allele of *cla-1.* First, *ola285* contains a missense mutation in the *cla-1* gene that converts Isoleucine (I) to Asparagine (N) at the residue 5753 (I5753N) (Figure 1K). Second, an independent allele of *clarinet*, *cla-1(ok560)*, phenocopied the ATG-9 localization defects observed for *ola285* mutants, both in terms of penetrance and expressivity (Figure 1L and M). Third, transheterozygous animals carrying both alleles *ola285* and *cla-1(ok560)* resulted in abnormal ATG-9 localization at synapses, similar to that seen for either *ola285* or *cla-1(ok560)* homozygous mutants (Figure 1M). The inability of *cla-1(ok560)* to complement the newly isolated allele *ola285* suggests that they correspond to genetic lesions within the same gene, *cla-1*. Together, our data indicate that CLA-1, an active zone protein with similarity to Piccolo and RIM, is required for trafficking of the autophagy protein ATG-9 at presynaptic sites.

### Clarinet long isoform, CLA-1L, acts cell autonomously to selectively regulate ATG-9 trafficking at presynaptic sites

The *cla-1* gene encodes three isoforms: CLA-1L (long), CLA-1M (medium) and CLA-1S (short) (Figure S1A). The long isoform, which contains a repetitive region predicted to be disordered (Figure 1K), is necessary for synaptic vesicle clustering, whereas the shorter isoforms are required for active zone assembly^40^. The isolated allele *cla-1*(*ola285)*, as well as the examined allele *cla-1(ok560)*, only affect CLA-1L, but not CLA-1M or CLA-1S. A null allele affecting all isoforms, *cla-1*(*wy1048*), did not display a more severe phenotype than the alleles affecting only CLA-1L (Figure S1A-B).

To determine the specific requirement of CLA-1L for ATG-9 trafficking at presynaptic sites in AIY, we manipulated the expression of CLA-1L in AIY using a cell-specific knockout strategy^40^. Briefly, the CLA-1L protein is characterized by having an unusually long and unique N-terminal region. We used a strain in which loxP sites were inserted, via CRISPR, to flank this unique region (Figure 1K; Figure S1A). Cell-specific expression of Cre recombinase in AIY, which leads to AIY-specific deletion of the CLA-1L isoform (without affecting CLA-1S and CLA-1M), resulted in the ATG-9 phenotype in AIY (Figure 1P and R), which is indistinguishable from that seen for the *cla-1* (*ok560)* allele (Figure 1Q and R, compare to wild type in Figure N, O and R). Together, our data indicate that the long isoform of CLA-1 (CLA-1L) regulates presynaptic trafficking of ATG-9 in a cell-autonomous manner.

Different CLA-1 isoforms play distinct roles in synaptic development, vesicle clustering and release^40^. While the *cla-1* phenotypes vary slightly between neurons, previous work demonstrated that the *cla-1* null allele, and not the alleles of the CLA-1 long isoform, displayed phenotypes on active zone composition in most neurons^40^. Consistent with these previous findings, we did not observe phenotypes for SYD-2/Liprin *α* when comparing its distribution in the presynaptic regions of AIY for wild-type and the *cla-1(ok560)* allele that specifically affects CLA-1L isoform (Figure S1 C-E). Based on the findings that 1) the *cla-1* null allele, but not *cla-1(L)*, displayed phenotypes on active zone composition, and 2) the *cla-1* null allele did not display a more severe ATG-9 phenotype than *cla-1(L)* mutants, we favor the hypothesis that the impaired ATG-9 trafficking is not due to defects in active zone assembly.

ATG-9 is a transmembrane protein transported to the presynaptic sites in vesicular structures^12, 21^. We therefore compared the localization of ATG-9::GFP with the synaptic vesicle integral membrane protein SNG-1/synaptogyrin1::BFP and the synaptic vesicle associated protein mCherry::RAB-3, both in wild-type animals and *cla-1(ola285)* mutants. In wild-type animals, we observed a strong colocalization between SNG-1 and RAB-3, as expected for synaptic vesicle proteins. ATG-9 also localized to presynaptic regions, largely colocalizing with these synaptic vesicle markers (Figure 2A-D). Interestingly, the phenotypes for synaptic vesicle proteins and ATG-9 differ in *cla-1(ola285)* mutants. Unlike the ATG-9::GFP foci observed in *cla-1(ola285)* mutants, the synaptic vesicle markers SNG-1 (or RAB-3) retained their wild type phenotype in *cla-1(ola285)* mutants in the Zone 2 region of AIY (Figure 2E-J). Our findings suggest that the long isoform of the active zone protein Clarinet specifically regulates ATG-9 trafficking at the presynaptic site, and that this trafficking is distinct from that of synaptic vesicle proteins.

**Figure 2.**
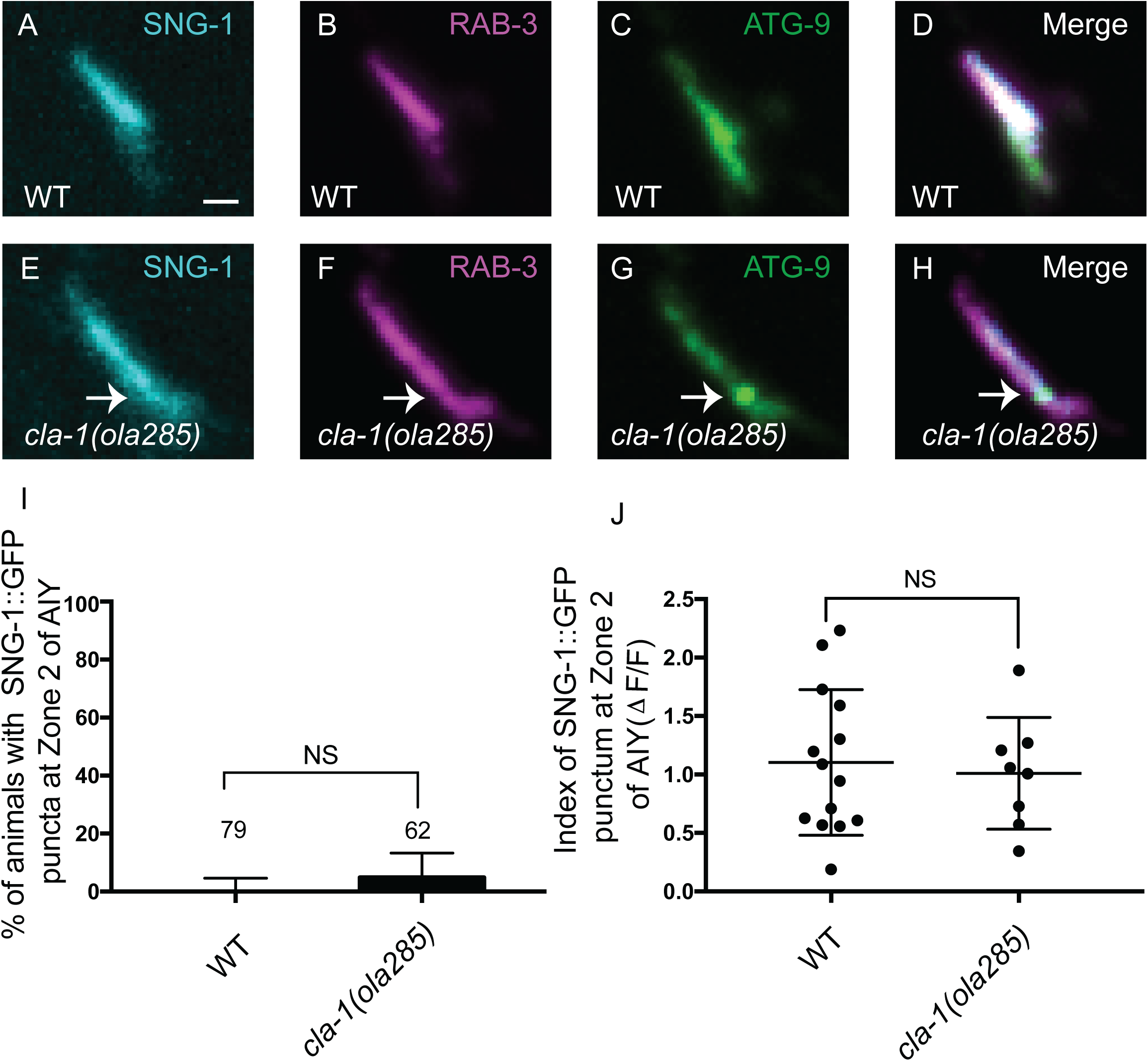
ATG-9 and synaptic vesicle proteins are differentially regulated by CLA-1L. (A-H) Distribution of SNG-1::BFP (pseudo-colored cyan) (A and E), mCherry::RAB-3 (pseudo-colored magenta) (B and F) and ATG-9::GFP (C and G) at Zone 2 of AIY (merge in D and H) in wild type (WT) (A-D) and *cla-1(ola285)* mutant (E-H) animals. While we observe a phenotype for abnormal ATG-9 distribution to subsynaptic foci in *cla-1(ola285)* mutants (indicated by arrows in E-H), we do not see a similar redistribution for synaptic vesicle proteins SNG-1 and RAB-3. (I) Quantification of the percentage of animals displaying SNG-1::GFP subsynaptic foci at AIY Zone 2 in different genotypes. Error bars represent 95% confidence interval. “NS” (not significant) by two-tailed Fisher’s exact test. The number on the bars indicates the number of animals scored. (J) Quantification of the index of SNG-1::GFP punctum (ΔF/F) at Zone 2 of AIY in wild type (WT) and *cla-1(ola285)* mutants. Error bars show standard deviation (SD). “NS” (not significant) by two-tailed unpaired Student’s t-test. Each dot in the scatter plot represents a single animal. Scale bar (in A for A-H), 1 μm.

### ATG-9 is enriched at endocytic intermediates in *cla-1(L)* mutants

ATG-9 undergoes exo-endocytosis at presynaptic sites, using synaptic vesicle cycling machinery^21^. The *cla-1(L)* phenotype is reminiscent of that seen for synaptic vesicle endocytosis mutants such as *unc-26/synaptojanin 1*, in which ATG-9 accumulates in purported endocytic intermediates, and abnormally colocalizes with the clathrin heavy chain subunit, CHC-1 (Figure S2A-E)^21^. To determine if, in *cla-1(L)* mutants, ATG-9 similarly abnormally accumulates in clathrin-rich endocytic intermediates, we examined the colocalization between ATG-9 and CHC-1 in wild type and *cla-1(ola285)* mutant animals. We observed that both ATG-9 and CHC-1 abnormally localized to synaptic foci in *cla-1(ola285)* mutants, and colocalized with each other (Figure 3A-H and Figure S2F).

**Figure 3.**
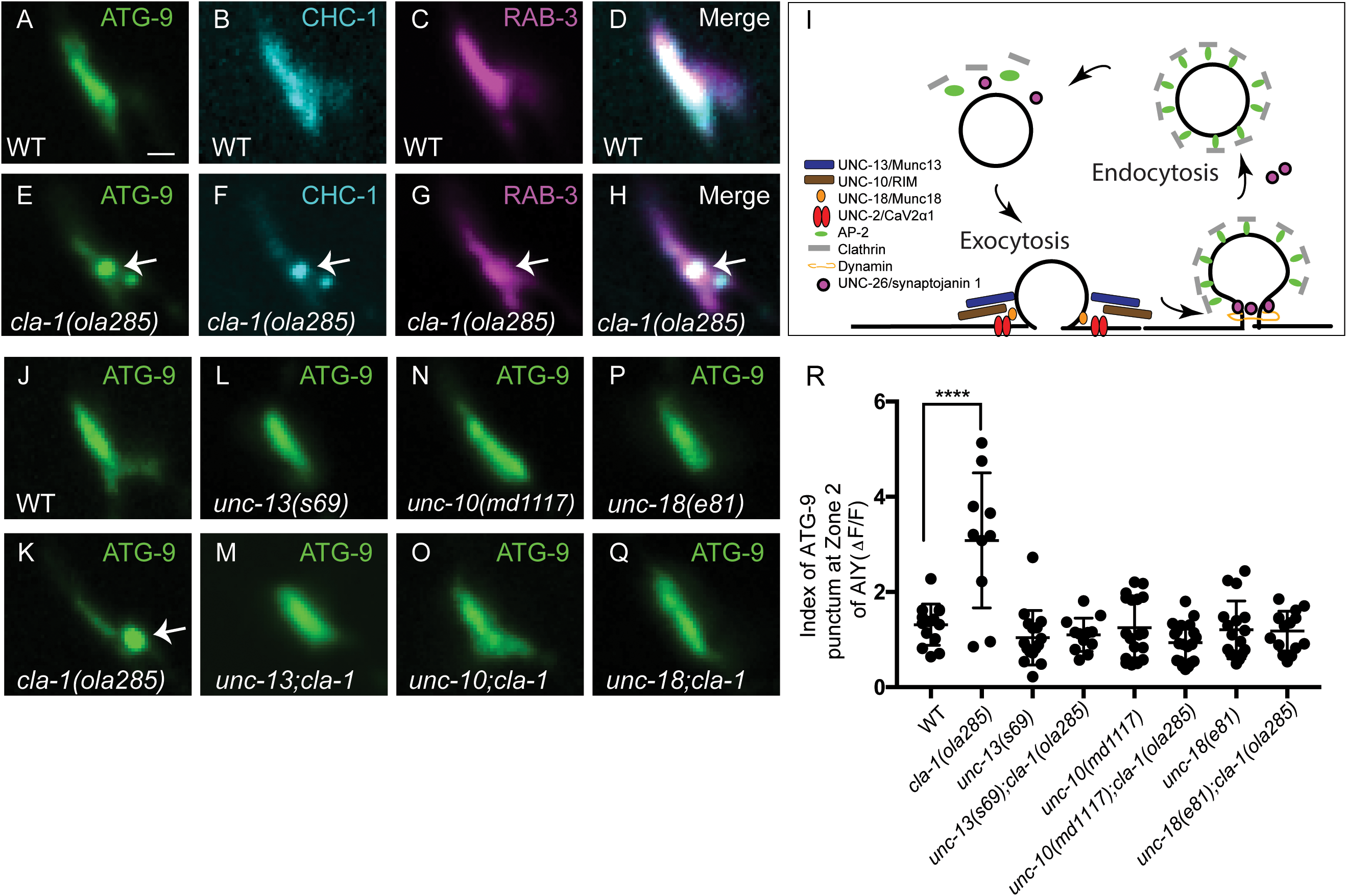
ATG-9 accumulates at clathrin-rich foci in *cla-1(ola285)* mutants and is suppressed by mutants for synaptic vesicle exocytosis. (A-H) Distribution of ATG-9::GFP (A and E), BFP::CHC-1 (pseudo-colored cyan) (B and F) and mCherry::RAB-3 (pseudo-colored magenta) (C and G) at Zone 2 of AIY (merge in D and H) in wild type (WT) (A-D) and *cla-1(ola285)* mutant (E-H) animals. ATG-9 subsynaptic foci are enriched with CHC-1 in *cla-1(ola285)* mutants (indicated by arrows in E-H). (I) Schematic of the proteins required for the synaptic vesicle cycle and associated with this study (both the names used for *C. elegans* and vertebrates are listed). (J-Q) Distribution of ATG-9::GFP at Zone 2 of AIY in wild type (WT) (J), *cla-1(ola285)* (K), *unc-13(s69)* (L), *unc-13(s69);cla-1(ola285)* (M), *unc-10 (md1117)* (N); *unc-10(md1117);cla-1(ola285)* (O), *unc-18(e81)* (P) and *unc-18(e81);cla-1(ola285)* (Q) animals. ATG-9 subsynaptic foci are indicated by the arrow (in K). (R) Quantification of the index of ATG-9 punctum (ΔF/F) at Zone 2 of AIY in the indicated genotypes. Error bars show standard deviation (SD). ****p<0.0001 by ordinary one-way ANOVA with Tukey’s multiple comparisons test. Each dot in the scatter plot represents a single animal. Scale bar (in A for A-H and J-Q), 1 μm.

To determine if the abnormal accumulation of ATG-9 in *cla-1(L)* mutants results from defects in ATG-9 sorting during exo-endocytosis, we next examined the necessity of synaptic vesicle exocytosis proteins in the ATG-9 phenotype of *cla-1(L)* mutants. We first visualized ATG-9 in putative null alleles *unc-13(s69)/Munc13, unc-10(md1117)/RIM, unc-18(e81)/Munc18,* and *unc-2(e55)/CaV2α1* (voltage-gated calcium channels), all of which encode proteins essential for synaptic vesicle fusion (Figure 3I)^48–55^. Single mutants of *unc-13(s69)*, *unc-10(md1117)*, *unc-18(e81)/Munc18* and *unc-2(e55)* did not disrupt ATG-9 localization (Figure 3L, N, P, R and Figure S2G). Double mutants of *unc-13(s69);cla-1(ola285)*, *unc-10(md1117);cla-1(ola285)*, *unc-18(e81);cla-1(ola285)* and *unc-2(e55);cla-1(ola285)* completely suppressed abnormal ATG-9 localization in *cla-1(L)* mutants (Figure 3M, O, Q, R and Figure S2G1). These results demonstrate that exocytosis is necessary for abnormal ATG-9 subsynaptic accumulation into clathrin-rich foci in *cla-1(ola285)* mutants, and support a model whereby the ATG-9 phenotype in *cla-1(L)* mutants results from defects in ATG-9 sorting during ATG-9 exo-endocytosis.

### CLA-1L extends to the periactive zone and genetically interacts with endocytic proteins to regulate ATG-9 trafficking

Presynapses have functional subdomains that coordinate distinct steps of the synaptic vesicle cycle: exocytosis takes place primarily at the active zone, while endocytosis takes place primarily at the periactive zone that surrounds the active zone^56–58^. Large active zone proteins which bear functional similarity to CLA-1L, such as Piccolo and Bassoon, extend across these subdomains and have been implicated in coupling synaptic vesicle exo-endocytosis at presynaptic sites in the sorting of synaptic components^57, 59–65^. CLA1L is twice the size as Piccolo and Bassoon, and contains largely disordered regions which could facilitate its extension from the active zone to the neighboring periactive zone. Consistent with this, in previous studies we documented that while the C-terminus of CLA-1 localized specifically to the active zone, the unique N-terminus of CLA-1L isoform localized beyond the active zone subdomain^40^.

To better examine the relationship between CLA-1L and the presynaptic subdomains, we compared the endogenous C-terminally tagged CLA-1::GFP, or the endogenous N-terminally tagged GFP::CLA-1L (Figure S3A-B)^40^, with a periactive zone marker APT-4/APA-2/AP-2α^66^. While the C-terminally tagged CLA-1::GFP specifically localizes to small puncta corresponding to the active zone, the N-terminally tagged GFP::CLA-1L displays a more distributed presynaptic pattern, extending to other regions of the synaptic bouton beyond the active zone (Figure 4A, D, H and K). Importantly, we observed that the N-terminally tagged GFP::CLA-1L, but not the C-terminally tagged CLA-1::GFP, colocalizes with the endocytic marker APT-4/APA-2/AP-2α (Figure 4A-N), which localizes to the periactive zones. These findings suggest that the long isoform of CLA-1 is anchored, via its C-terminus, to the active zone, but extends to the periactive zone where endocytosis occurs.

**Figure 4.**
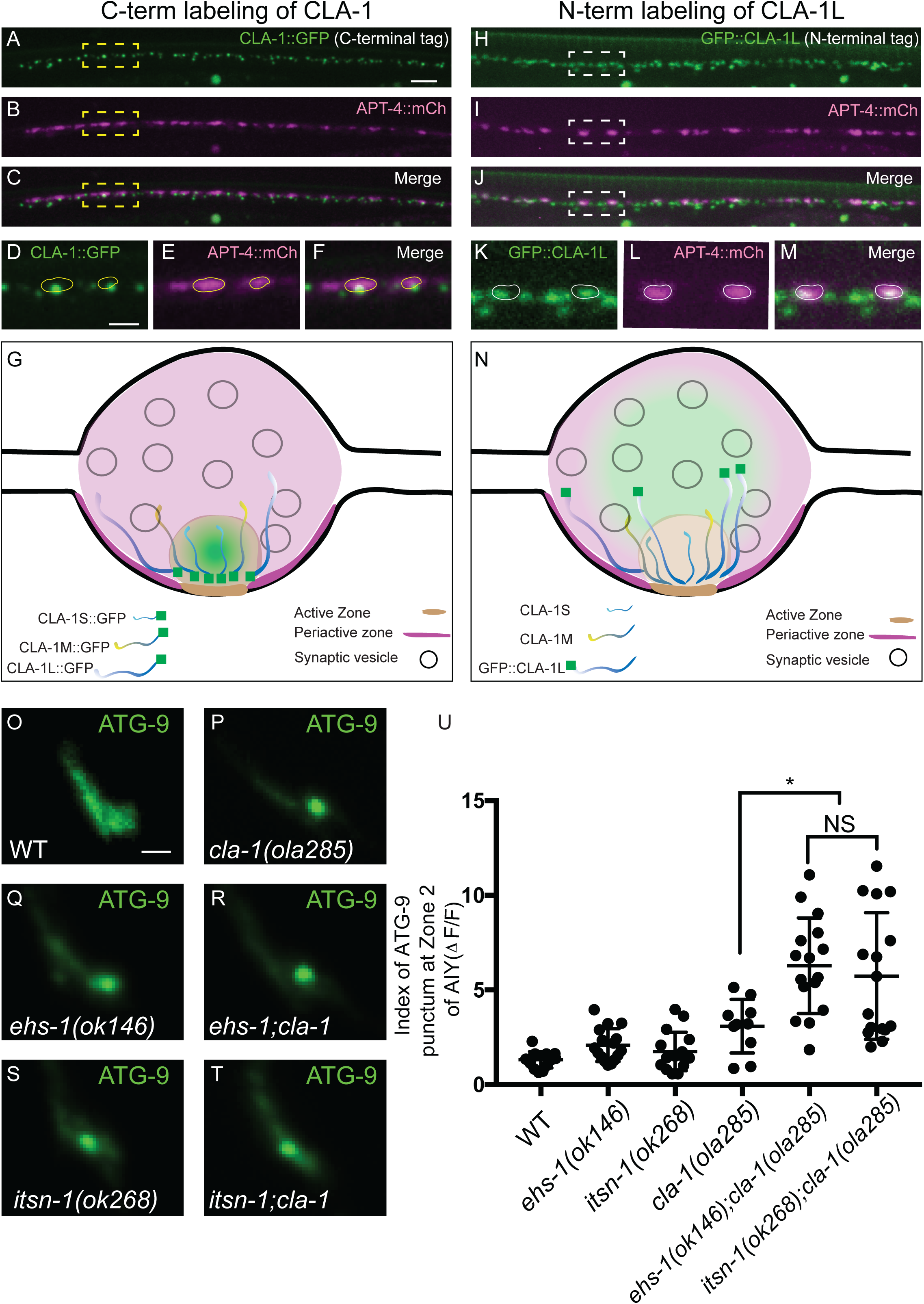
CLA-1L genetically interacts with endocytic proteins at the periactive zone to regulate ATG-9 trafficking. (A-C) Distribution of endogenous C-terminally tagged CLA-1::GFP (A) and the endocytic zone marker APT-4/APA-2/AP-2α::mCherry (APT-4::mCh, pseudo-colored magenta) (B) in the neurons of the posterior dorsal nerve cord (merge in C) in wild type animals. (D-F) Enlarged regions enclosed in dashed boxes in A-C. Endogenous C-terminally tagged CLA-1::GFP (D) localizes to small puncta corresponding to the active zone^40^, and different from the pattern observed for periactive zone protein, APT-4::mCh (E, merge in F). Yellow circles are drawn based on the outline of APT-4::mCh puncta in E, and are located at the same positions in D-F. (G) Schematic of the localization of the C-terminally tagged CLA-1::GFP, relative to the subsynaptic active and periactive zones. (H-J) Distribution of endogenous N-terminally tagged GFP::CLA-1L (H) and the endocytic zone marker APT-4/APA-2/AP-2α::mCherry (APT-4::mCh, pseudo-colored magenta) (I) in neurons of the posterior dorsal nerve cord (merge in J) in wild type animals. (K-M) Enlarged regions enclosed in dashed boxes in H-J. Endogenous N-terminally tagged GFP::CLA-1L (K) displays a more distributed synaptic distribution as compared to the C-terminally tagged CLA-1:GFP (compare with A, D and F, see also ^40^) and colocalizes with APT-4::mCh (L, merge in M). White circles are drawn based on the outline of APT-4::mCh puncta in L, and are located at the same positions in K-M. (N) Schematic of the localization of the N-terminally tagged GFP::CLA-1L, relative to the subsynaptic active and periactive zones. (O-T) Distribution of ATG-9::GFP at Zone 2 of AIY in wild type (WT) (O), *cla-1(ola285)* (P), *ehs-1(ok146)* (Q), *ehs-1(ok146);cla-1(ola285)* (R), *itsn-1(ok268)* (S) and *itsn-1(ok268);cla-1(ola285)* (T) mutant animals. (U) Quantification of the index of ATG-9 punctum (ΔF/F) at Zone 2 of AIY in the indicated genotypes. Error bars show standard deviation (SD). “NS” (not significant), *p<0.05 by ordinary one-way ANOVA with Tukey’s multiple comparisons test. Each dot in the scatter plot represents a single animal. Scale bar (in A for A-C and H-J), 5 μm; (in D for D-F and K-M), 2 μm; (in O for O-T), 1 μm.

Based on the localization of CLA-1L to these presynaptic subdomains, and the ATG-9 phenotypes observed in *cla-1(L)* and endocytic mutants, we hypothesized the existence of genetic interactions between CLA-1L and endocytic proteins at the periactive zone. To test this hypothesis, we examined the localization of ATG-9 in the double mutants of *cla-1(ola285)* with genes encoding periactive zone endocytic proteins, *ehs-1(ok146)/EPS15* or *itsn-1(ok268)/intersectin 1*. We focused our analyses on the endocytic regulators EHS-1/EPS15 and ITSN-1/intersectin 1 because of their hypothesized roles in coupling synaptic vesicle exocytosis at the active zone, and endocytosis at the periactive zone^56, 57, 67–74^. We observed that in null alleles of *ehs-1(ok146)/EPS15* and *itsn-1(ok268)/intersectin 1*, thirty percent of worms displayed abnormal ATG-9 foci (compared to sixty percent in *cla-1(L)* mutants) (Figure 4Q, S and Figure S3C). Interestingly, we observed that *ehs-1(ok146);cla-1(ola285)* and *itsn-1(ok268)*;*cla-1(ola285)* enhanced the ATG-9 phenotype as compared to any of the single mutants (Figure 4O-T), both in terms of penetrance (Figure S3C) and expressivity (Figure 4U). Our findings uncover a cooperative genetic relationship between CLA-1L and the EHS-1-ITSN-1 endocytic scaffolding complex, suggesting that the active zone protein CLA-1L acts in pathways that are partially redundant to the EHS-1-ITSN-1 complex in linking the active zone and periactive zone regions to regulate ATG-9 trafficking at presynapses.

### The clathrin adaptor complexes, AP-2 and AP180, regulate ATG-9 trafficking at presynaptic sites

Endocytosis and sorting of transmembrane proteins depend on adaptor protein complexes such as AP-2^75–77^. Given our observations that in *cla-1(ola285)* mutants, ATG-9 abnormally accumulates in clathrin-rich endocytic intermediates, and that the N-terminal region of CLA-1L co-localizes with AP-2 in the periactive zone, we next examined the genetic relationship between clathrin adaptor protein complexes and CLA-1L in ATG-9 localization. The AP-2 complex mediates clathrin-mediated endocytosis (CME) of synaptic vesicle proteins^78–83^, and it has been implicated in the sorting of ATG-9 during autophagy induction in mammalian non-neuronal cells^84–86^. To determine if the AP-2, and the associated AP180, adaptor complexes were required in presynaptic trafficking of ATG-9, we examined ATG-9 localization in the null alleles *dpy-23(e840)/AP2µ* and *unc-11(e47)/AP180*. We observed that *dpy-23(e840)/AP2µ* and *unc-11(e47)/AP180* mutants phenocopied *cla-1(ola285)* mutants in ATG-9 presynaptic trafficking defects (Figure 5A, B, C, E and F). In addition, the expressivity of the ATG-9 trafficking defects was enhanced in *dpy-23(e840)/AP2µ;cla-1(ola285)* double mutant worms (Figure 5D-F). These findings suggest shared mechanisms that similarly result in defective ATG-9 trafficking when clathrin-associated adaptor complexes, or the active zone protein CLA-1L, are disrupted.

**Figure 5.**
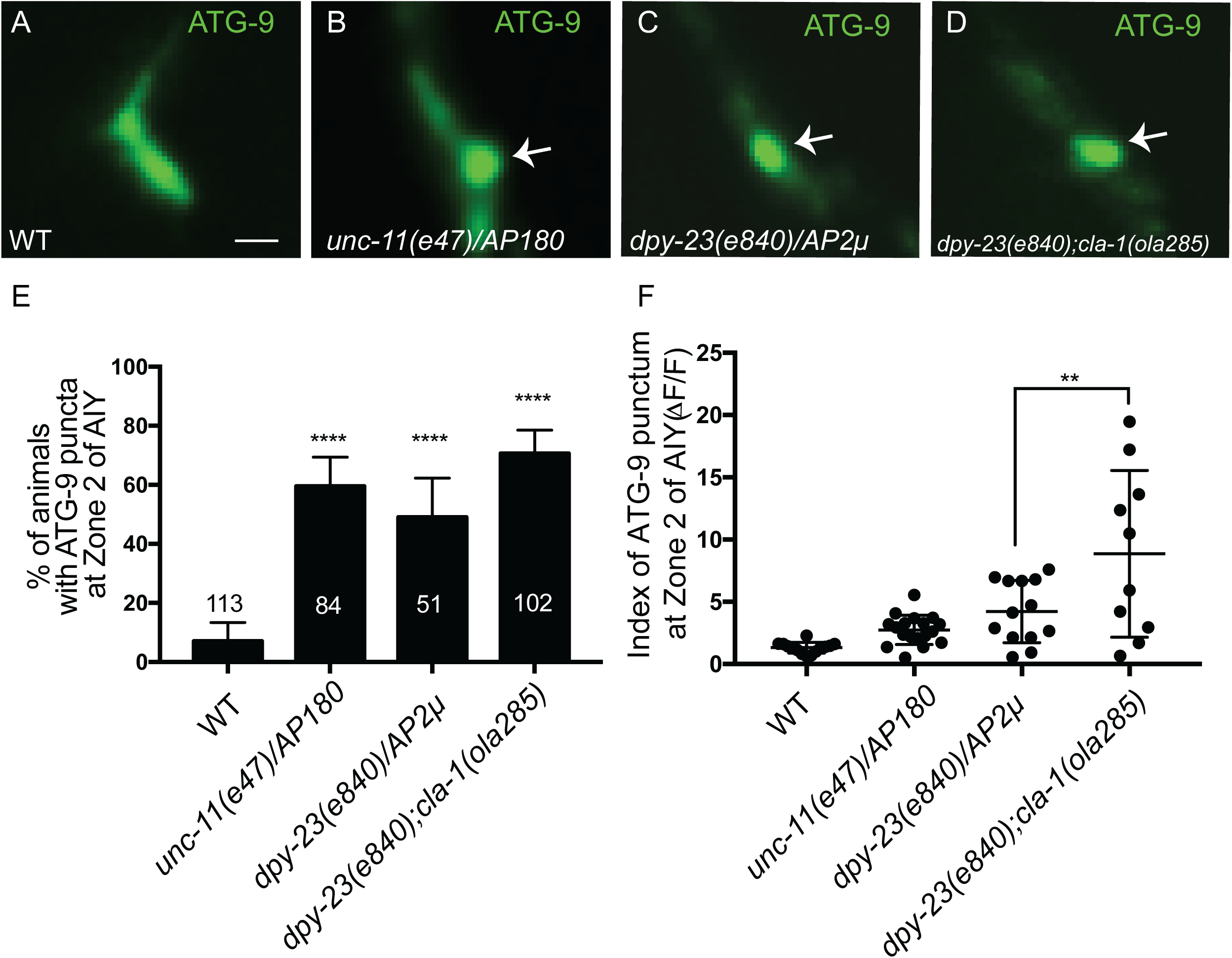
The clathrin-associated adaptor complexes, AP-2 and AP180, regulate ATG-9 trafficking at presynaptic sites. (A-D) Distribution of ATG-9::GFP at Zone 2 of AIY in wild type (WT) (A), *unc-11(e47)/AP180* (B), *dpy-23(e840)/AP2µ* (C) and *dpy-23(e840);cla-1(ola285)* (D) mutant animals. Abnormal ATG-9 subsynaptic foci are indicated by arrows in B-D. (E) Quantification of the percentage of animals displaying ATG-9 subsynaptic foci at AIY Zone 2 for the indicated genotypes. Error bars represent 95% confidence interval. ****p<0.0001 by two-tailed Fisher’s exact test. The number on the bars indicates the number of animals scored. (F) Quantification of the index of ATG-9 punctum (ΔF/F) at Zone 2 of AIY for the indicated genotypes. Error bars show standard deviation (SD). **p<0.01 by ordinary one-way ANOVA with Tukey’s multiple comparisons test. Each dot in the scatter plot represents a single animal. Scale bar (in A for A-D), 1 μm.

### ATG-9 is sorted to endocytic intermediates via SDPN-1/syndapin 1 and the AP-1 adaptor complex

Clathrin-associated adaptor complexes mediate internalization and sorting of cargoes from both the plasma membrane and intracellular organelles^87, 88^. Given the distinct phenotypes observed for synaptic vesicle proteins and ATG-9 upon disruption of CLA-1L (Figure 2), we next investigated pathways associated with clathrin-dependent endosomal sorting, the disruption of which could lead to the abnormal accumulation of ATG-9 in endocytic intermediates at presynaptic sites.

We first examined SDPN-1/syndapin 1, a protein known to play important roles in early stages of membrane invagination during both activity-dependent bulk endocytosis (ADBE)^89, 90^ and ultrafast endocytosis^91^. We reasoned that if ATG-9 was sorted via SDPN-1-dependent mechanisms, then *sdpn-1* mutants would suppress the observed ATG-9 foci for *cla-1(L)* mutants and mutants for the clathrin-associated adaptor complexes. We observed that while ATG-9 localization was not disrupted in *sdpn-1(ok1667)* single mutants, the abnormal ATG-9 foci were suppressed in *sdpn-1(ok1667);cla-1(ola285)* or *sdpn-1(ok1667);unc-11(e47)/AP180* double mutant animals (Figure 6A-H). These findings are consistent with the requirement of SDNP-1 in the transport of ATG-9 to intermediate endosomal structures, from which CLA-1L and the clathrin-associated adaptor complexes sort.

**Figure 6.**
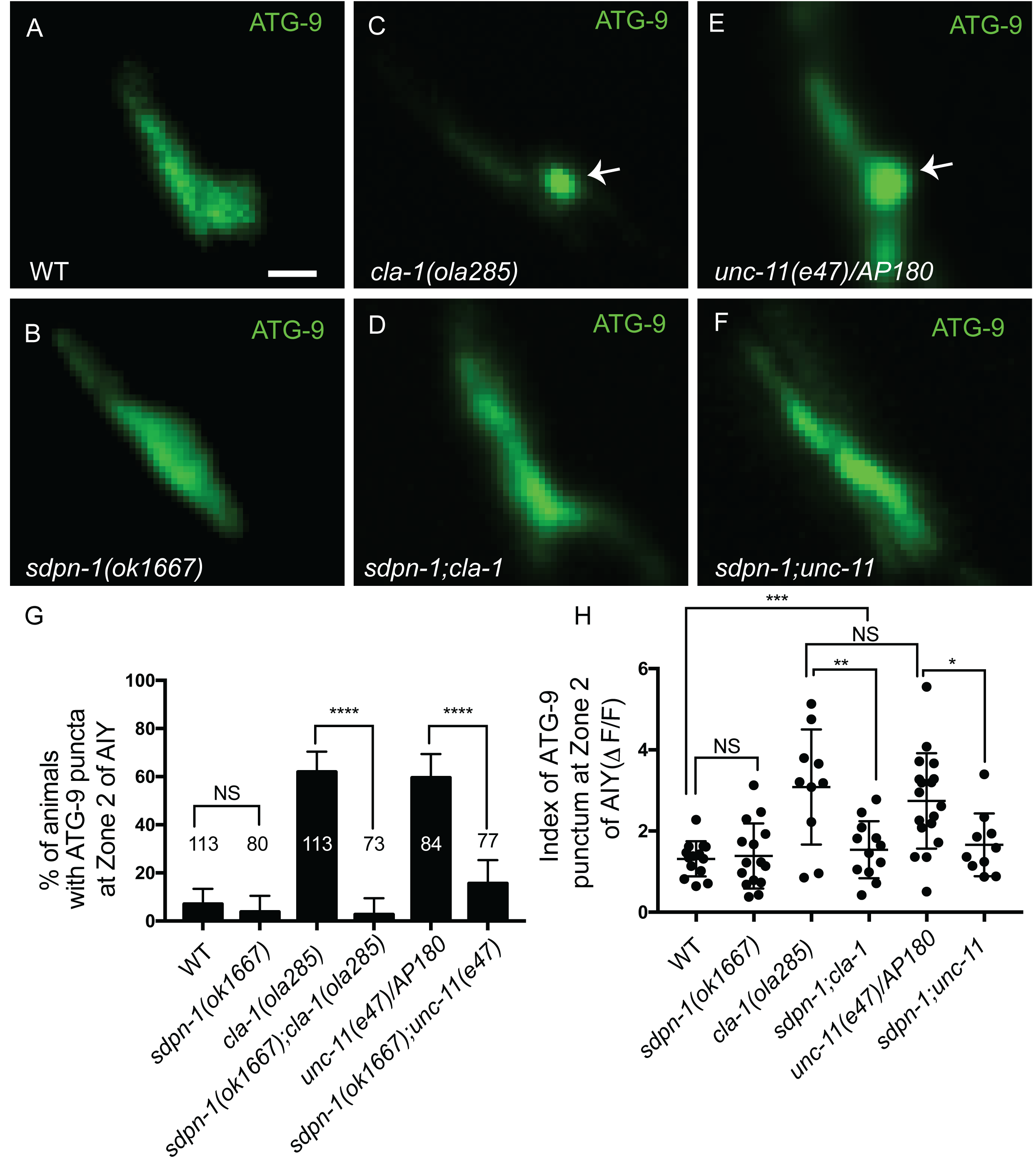
SDPN-1/syndapin 1 regulates ATG-9 sorting at presynaptic sites. (A-F) Distribution of ATG-9::GFP at Zone 2 of AIY in wild type (WT) (A), *sdpn-1(ok1667)* (B), *cla-1(ola285)* (C), *sdpn-1(ok1667);cla-1(ola285)* (D), *unc-11(e47)/AP180* (E) and *sdpn-1(ok1667);unc-11(e47)* (F) mutant animals. Abnormal ATG-9 subsynaptic foci are indicated by arrows in C and E. Note that mutations in SDPN-1/syndapin 1 suppress the abnormal ATG-9 phenotypes in *cla-1* and *unc-11/AP180* mutants. (G) Quantification of the percentage of animals displaying abnormal ATG-9 subsynaptic foci at AIY Zone 2 for the indicated genotypes. Error bars represent 95% confidence interval. “NS” (not significant), ****p<0.0001 by two-tailed Fisher’s exact test. The number on the bars indicates the number of animals scored. (H) Quantification of the index of ATG-9 punctum (ΔF/F) at Zone 2 of AIY for the indicated genotypes. Error bars show standard deviation (SD). “NS” (not significant), *p<0.05, **p<0.01, ***p<0.001 by ordinary one-way ANOVA with Tukey’s multiple comparisons test. Each dot in the scatter plot represents a single animal. Scale bar (in A for A-F), 1 μm.

Next, we examined the AP-1 adaptor complex, which acts at presynaptic sites to mediate endosomal sorting of activity-dependent bulk endocytosis (ADBE)^92–94^. To determine the requirement of the AP-1 adaptor complex in ATG-9 sorting at presynaptic sites, we examined ATG-9 localization in *unc-101(m1)/AP1µ1* single mutant animals and in double mutants with *cla-1(ola285)*, *dpy-23(e840)/AP2µ* and *unc-11(e47)/AP180*. We observed that while *unc-101(m1)/AP1µ1* single mutant animals did not display detectable phenotypes in ATG-9 localization (Figure 7C, I and Figure S4F), *unc-101(m1)/AP1µ1* suppressed the ATG-9 phenotype in double mutant animals (Figure 7A-D, G-I and Figure S4B, D, E and F).

**Figure 7.**
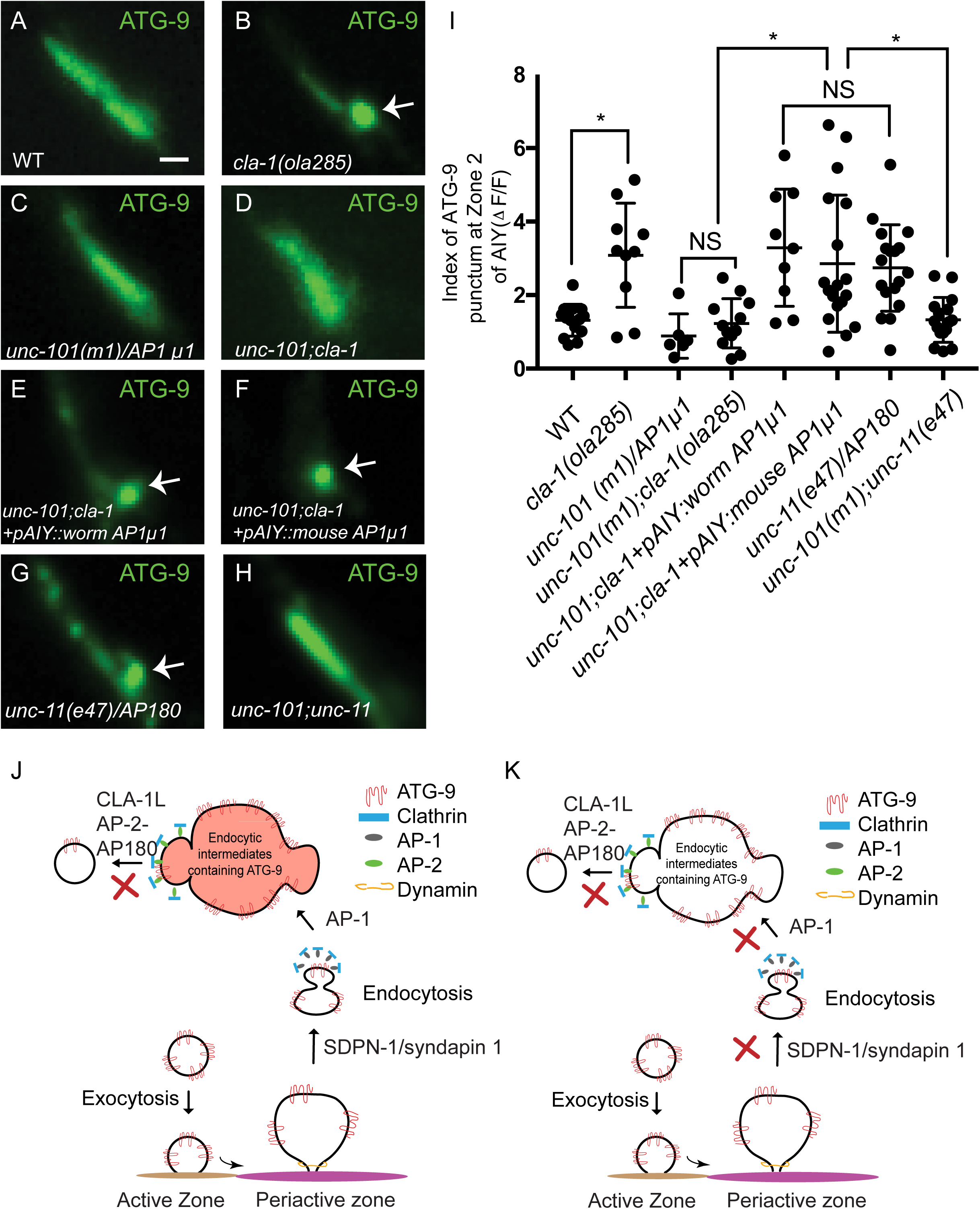
ATG-9 is sorted to the endocytic intermediates via the AP-1 adaptor complex. (A-H) Distribution of ATG-9::GFP at Zone 2 of AIY in wild type (WT) (A), *cla-1(ola285)* (B), *unc-101(m1)/AP-1µ1* (C), *unc-101(m1);cla-1(ola285)* (D), *unc-101;cla-1* mutants with *C. elegans* UNC-101/AP-1µ1 cDNA cell-specifically expressed in AIY (E), *unc-101;cla-1* mutants with mouse AP1µ1 cDNA cell-specifically expressed in AIY (F), *unc-11(e47)/AP180* (G) and *unc-101(m1);unc-11(e47)* (H). Abnormal ATG-9 subsynaptic foci are indicated by arrows in B and E-G. (I) Quantification of the index of ATG-9 punctum (ΔF/F) at Zone 2 of AIY for indicated genotypes. Error bars show standard deviation (SD). “NS” (not significant), *p<0.05 by ordinary one-way ANOVA with Tukey’s multiple comparisons test. Each dot in the scatter plot represents a single animal. (J) Cartoon diagram of the genetic interactions, and model, in this study. Mutants for CLA-1L, AP-2 and AP180 adaptor complexes display similar ATG-9 phenotypes at synapses, and are necessary for resolving ATG-9-containing foci (clathrin rich endocytic intermediates). (K) Cartoon diagram of the genetic interactions, and model displaying the genetic interactions of CLA-1L with SDPN-1/syndapin 1 and the AP-1 adaptor complex, which suppress the ATG-9 phenotypes observed for *cla-1(L)*, *AP-2* or *AP180* mutants, consistent with roles for SDPN-1/syndapin 1 and the AP-1 upstream of the formation of the abnormal ATG-9 foci (similar to what was observed for exocytosis mutants in Figure 3, which also suppressed ATG-9 phenotypes in *cla-1(L)* mutants). Scale bar (in A for A-H), 1 μm.

To confirm this, we made double mutants of *cla-1(ola285)* with another allele, *unc-101(sy108)*, which also suppressed the *cla-1(ola285)* ATG-9 phenotype (Figure S4F). Furthermore, single-cell expression of the *C. elegans* cDNA of *unc-101(m1)/AP1µ1* in AIY in the *unc-101(m1);cla-1(ola285)* double mutants reverted the phenotype, indicating that AP-1 acts cell autonomously in AIY to suppress the *cla-1(L)* phenotype (Figure 7E, I and Figure S4F). The *C. elegans* UNC-101/AP1µ1 is more similar to the murine AP1µ1 (Query Cover: 100%; Percentage Identity: 74%) than to the murine AP2µ (Query Cover: 98%; Percentage Identity: 41%) (Figure S4A). To examine the conserved role of UNC-101/AP1µ1 in ATG-9 trafficking, we expressed murine cDNA of *AP1µ1* or *AP2µ* cell specifically in AIY of the *unc-101(m1);cla-1(ola285)* double mutants, and examined ATG-9 localization. We observed that expression of murine *AP1µ1*, but not *AP2µ* cDNA, reverted the suppression seen for *unc-101(m1);cla-1(ola285)* double mutant animals (Figure 7F, I and Figure S4C and F). Our data indicate that the AP-1 complex regulates ATG-9 presynaptic trafficking cell-autonomously in AIY and that this function of AP-1 is conserved between mouse and *C. elegans*. Importantly, these findings indicate that the AP-1 complex mediates ATG-9 trafficking to the subsynaptic foci in *cla-1(L)* mutants, in *AP-2* mutants and in *AP180* mutants. Together, our findings are consistent with a model whereby ATG-9 is sorted to endocytic intermediates via proteins like AP-1 and SDPN-1, while CLA-1L, AP-2 and AP180 sort ATG-9 out of those endocytic intermediates (Figure 7J and K).

### Disrupted ATG-9 trafficking in *cla-1(L)* mutants is associated with a deficit in activity-induced autophagosome formation

ATG-9 is important for autophagosome biogenesis at synapses^12^. To examine how autophagosome formation is affected in *cla-1(L)* mutants, we measured the average number of LGG-1/Atg8/GABARAP puncta (an autophagosomal marker) in the AIY neurites^12, 15, 21^ (Figure 8A-C). Previously we observed the average number of LGG-1 puncta increased in AIY when the wild-type animals were cultivated at 25°C^15^, a condition known to increase the activity state of the AIY neurons^95^. Worms with impaired exocytosis (in *unc-13* mutants) or impaired autophagy (in *atg-9* mutants) failed to display increased LGG-1 synaptic puncta^15, 21^. We found that, unlike wild type animals, the average number of LGG-1 puncta did not increase in *cla-1(L)* mutants (alleles *ola285* and *ok560*) in response to cultivation temperatures that increase the activity state of the neuron (Figure 8D and S5A). These findings indicate that activity-induced autophagosome formation at synapses is impaired in *cla-1(L)* mutants.

**Figure 8.**
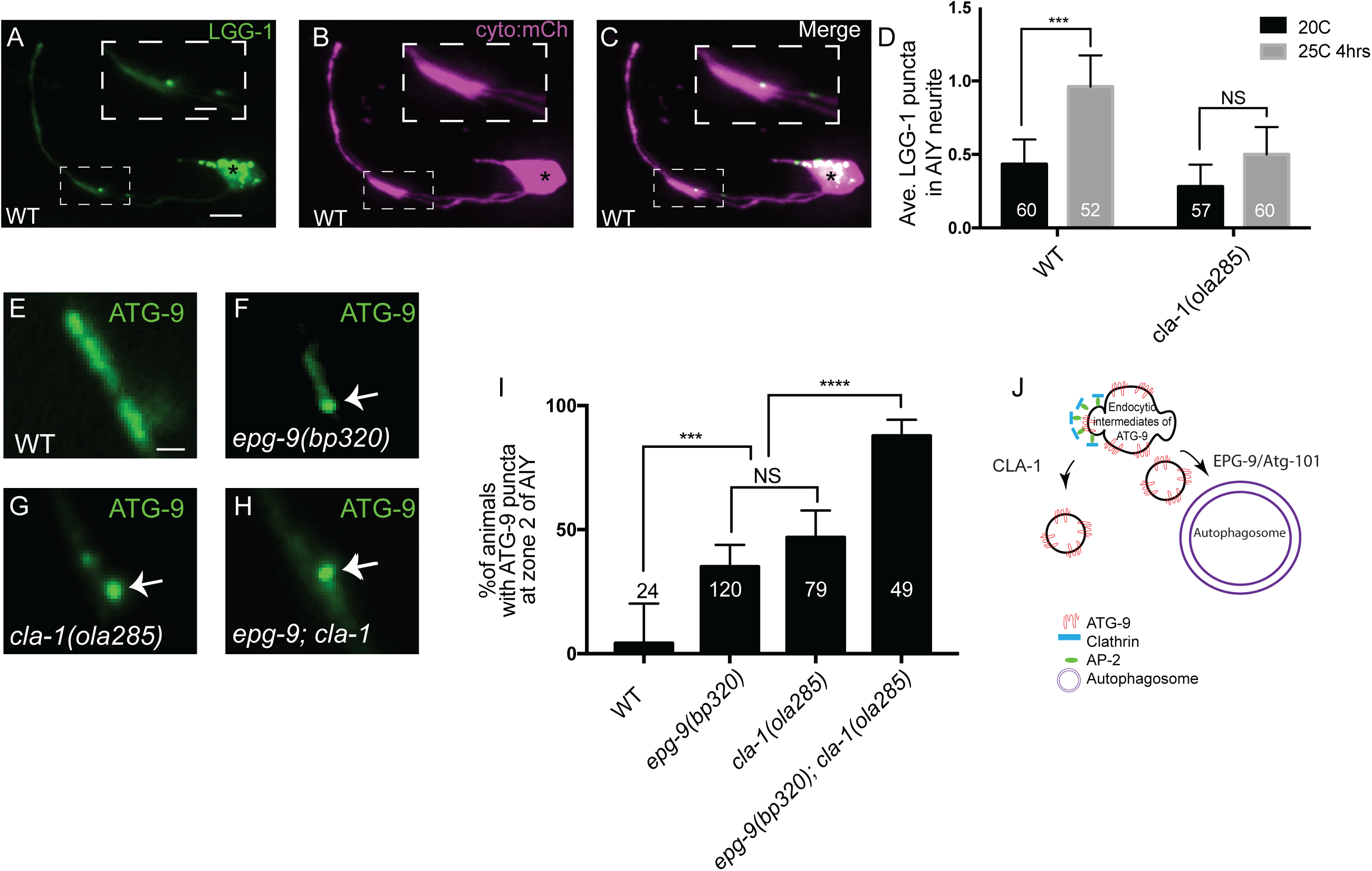
Disrupted ATG-9 trafficking in *cla-1(L)* mutants is associated with a deficit in activity-induced autophagosome formation. (A-C) Confocal micrographs of GFP::LGG-1 (A) and cytoplasmic mCherry (cyto::mCh) (pseudo-colored magenta, B) in AIY (merge in C). Inset is the enlarged region enclosed in dashed box to show one LGG-1 punctum in AIY synaptic Zone 2. (D) Quantification of the average number of LGG-1 puncta in the AIY neurites at 20°C and at 25°C for 4 hours in wild type (WT) and *cla-1(ola285)* mutants. (Note: The activity state of the thermotaxis interneurons AIY is reported to increase when animals are cultivated at 25°C for 4 hours, compared to animals at 20°C^95, 139, 140^. Error bars represent 95% confidence interval. “NS” (not significant), ***p<0.001 by Kruskal-Wallis test with Dunn’s multiple comparisons test. The number on the bars indicates the number of animals scored. (E-H) Distribution of ATG-9::GFP at Zone 2 of AIY in wild type (E), *epg-9(bp320)* (F), *cla-1(ola285)* (G) and *epg-9(bp320); cla-1(ola285)* (H) mutant animals. Arrows (in F-H) indicate abnormal ATG-9 foci. (I) Quantification of the percentage of animals displaying ATG-9 subsynaptic foci at AIY Zone 2 in the indicated genotypes. Error bars represent 95% confidence interval. “NS” (not significant), ***p<0.001, ****p<0.0001 by two-tailed Fisher’s exact test. The number on the bar indicates the number of animals scored. (J) Cartoon diagram representing autophagosome biogenesis at the synapse, and our model on its relationship with ATG-9 trafficking. The ATG-9 phenotype is enhanced in *epg-9(bp320); cla-1(ola285)* double mutants. Scale bar (in A for A-C), 5 μm; (in inset of A for inset of A-C), 2μm; (in E for E-H), 1 μm.

Previously, we found that in mutants that affect early stages of autophagy such as *epg-9(bp320)*, ATG-9 accumulated into subsynaptic foci, which colocalized with the clathrin heavy chain CHC-1^21^. To determine a potential cross-talk between CLA-1L-mediated ATG-9 endocytosis and autophagy, we generated *epg-9(bp320);cla-1(ola285)* double mutant animals. Both the penetrance and the expressivity of the ATG-9 phenotype are enhanced in the double mutants, compared to single mutants (Figure 8E-I and S5B). The observed enhancement suggests the existence of parallel pathways that sort ATG-9 at the synapse during exo-endocytosis and that lead to autophagosome formation (Figure 8J). Importantly, our data uncover a relationship between ATG-9 trafficking at the synapse, and the activity-dependent regulation of synaptic autophagy. Together, our findings are consistent with a model whereby disrupted ATG-9 trafficking in *cla-1(L)* mutants contributes to deficits in activity-induced autophagosome formation at synapses.

## Discussion

The long isoform of the active zone protein Clarinet (CLA-1L), regulates ATG-9 trafficking at synapses and presynaptic autophagy. Autophagy, a conserved cellular degradative pathway, is temporally and spatially regulated in neurons to occur at synaptic compartments and in response to increased synaptic activity states^6, 8, 10–15, 18, 24^. How synaptic autophagy and synaptic activity states are coordinated in neurons is not well understood. From unbiased forward genetic screens, we uncover a role for the active zone protein Clarinet in activity-dependent autophagosome formation at synapses, and in regulated synaptic trafficking of the autophagy protein, ATG-9. Our findings are consistent with recent studies in primary hippocampal neurons which demonstrated that active zone proteins such as Bassoon and Piccolo play important roles in synaptic protein homeostasis^96^, in part via the regulation of presynaptic autophagy^27–29^. Bassoon negatively regulates presynaptic autophagy by serving as a scaffold for Atg5^27^, and the E3 ubiquitin ligase Parkin^28, 29^. Our studies extend these findings *in vivo* in *C. elegans*, providing further evidence for the specific roles of active zone proteins in the temporal and spatial regulation of activity-dependent synaptic autophagy. However, the mechanisms of CLA-1L regulation of synaptic autophagy are distinct from those observed for Bassoon. We find that instead of inhibiting autophagy, CLA-1L is required for activity-dependent synaptic autophagy, likely via the regulated trafficking of ATG-9 at synapses.

CLA-1L regulates presynaptic sorting of ATG-9 during exo-endocytosis, and genetically interacts with the AP-1 and AP-2 adaptor complexes in mediating ATG-9 trafficking at presynaptic sites. ATG-9 is the only transmembrane protein in the core autophagy pathway, and plays key roles as a lipid scramblase to nucleate the growth of the autophagosome isolation membrane^97–100^. In both yeast and mammalian cells, ATG-9 trafficking is regulated, and its subcellular localization is associated with the promotion of local autophagosome formation^35–39, 84, 101–105^. Local autophagosome formation at synapses is associated with the degradation of local synaptic cargo^6, 15, 27, 106–111^, and ATG-9 trafficking at presynaptic sites via exo-endocytosis could link synaptic activity states with local synaptic autophagy^12, 21^. We find that the active zone protein CLA-1L regulates ATG-9 trafficking during its exo-endocytosis at presynaptic sites. In *cla-1(L)* mutants, ATG-9 abnormally accumulates at clathrin-rich presynaptic foci, suggesting abnormal localization of ATG-9 into endocytic intermediates. Furthermore, we find that CLA-1L genetically interacts with clathrin-associated adaptor complexes, such as AP-1, AP-2 and AP180, in mediating ATG-9 presynaptic trafficking. Our findings are consistent with studies in non-neuronal cells which demonstrated that the AP-1 and AP-2 complexes mediate ATG-9 trafficking between the plasma membrane, the trans-Golgi network (TGN), the recycling endosomes and the growing autophagosomes^84, 86, 112, 113^. Our findings are also consistent with recent studies that uncovered roles for synaptojanin 1/UNC-26-mediated ATG-9 exo-endocytosis at synapses^21^. Our data extend these studies and suggest a model in which the active zone protein CLA-1L is necessary for normal ATG-9 trafficking at synapses *in vivo*, likely via the regulated clathrin-dependent sorting of ATG-9 during exo-endocytosis.

CLA-1L spans from the exocytic active zone to the endocytic periactive zone and regulates ATG-9 sorting. Clarinet is an active zone protein involved in the localization of synaptic vesicle proteins, in active zone assembly, and in neurotransmission^40^. These roles of CLA-1 are genetically separable. For example, while null alleles of *cla-1* that affect all three isoforms disrupt active zone assembly and synapse number, alleles that specifically affect the CLA-1 long isoform (CLA-1L) display specific defects in synaptic vesicle clustering, without disrupting active zone assembly in most examined neurons (Figure S1)^40^. We similarly observe specific roles for the CLA-1L isoform in ATG-9 sorting, which we interpret as distinct from the other roles of CLA-1 in active zone assembly for two reasons. First, the *cla-1* null allele did not display a more severe ATG-9 phenotype than *cla-1(L)* mutants, suggesting that the ATG-9 sorting defects are due to the loss of CLA-1L. Second, we do not observe ATG-9 sorting phenotypes in mutants for other active zone proteins SYD-1/mSYD1A and SYD-2/Liprin-⍺, known to disrupt active zone assembly (Data not shown)^114–116^. The PDZ and C2 domains of CLA-1 have sequence homology to active zone proteins RIM, Fife and Piccolo, and CLA-1 has been proposed to be functionally similar to Piccolo and Bassoon in its roles at the active zone^40^. In vertebrate synapses, Piccolo and Bassoon extend from the active zone region to periactive zones, and this architecture has been proposed to be important in their purported functional roles coupling exocytosis at the active zone, and protein sorting during endocytosis at the periactive zone^57, 59–65^. CLA-1L is an 8922 amino-acid protein, twice the size of Bassoon (3942 amino acids) and Piccolo (4969 amino acids). It is anchored to the active zone via its C-terminus^40^, with a large disordered N-terminal domain extending to the periactive zone, where endocytic processes occur. Consistent with these observations on CLA-1L size and position at the synapses, we observe that CLA-1L regulates ATG-9 trafficking by genetically interacting with the periactive zone proteins EHS-1/EPS15 or ITSN-1/intersectin 1, which have been suggested as linkers between active-zone exocytosis and periactive zone endocytosis^56, 57, 67–74^. Our observation that the *ehs-1(ok146)* null allele^117^ and *itsn-1(ok268)* null allele^118^ display enhanced defects in ATG-9 sorting in *cla-1L (ola285)* mutants is consistent with a model of functional redundancy between these proteins and CLA-1L in ATG-9 sorting. We propose that the specific requirement of CLA-1L for sorting of ATG-9 at synapses is mediated via its capacity to extend across presynaptic sub-domains, from the exocytic active zone to the endocytic periactive zone.

ATG-9 likely undergoes endosome-mediated endocytosis at presynaptic sites. The synaptic vesicle cycle is supported by different modes of endocytosis^119, 120^, which include clathrin-mediated endocytosis (CME)^78–80, 121–123^, endosome-mediated endocytosis (activity-dependent bulk endocytosis (ADBE)^90, 92, 124–126^ and ultrafast endocytosis^127, 128^. In contrast to the better-known mechanisms underpinning recycling of synaptic vesicle proteins, less is known about the mechanisms of ATG-9 exo-endocytosis. While we find that ATG-9 presynaptic trafficking depends on proteins such as UNC-13/Munc13 and AP-2, which also regulate synaptic vesicle protein trafficking, the phenotypes of synaptic vesicle proteins and ATG-9 are distinct for *cla-1(L)* (Figure 2)*, AP-2* or *AP180* mutant animals (data not shown). These findings indicate that although ATG-9 and synaptic vesicle proteins use the same exo-endocytosis machinery, they are likely trafficked and sorted via different pathways. By examining the genetic interactions of adaptor protein complexes and endocytic factors in the regulation of ATG-9 trafficking, we propose a model whereby the AP-1 adaptor complex and the F-BAR protein syndapin I (SDPN-1) mediate trafficking of ATG-9 to a transient sorting station from which AP2-AP180 complexes facilitate clathrin-mediated ATG-9 vesicle budding (Figure 9). Consistent with observations in yeast^36–39^, *C. elegans*^12^ and non-neuronal mammalian cells^35^, our findings suggest that this regulated ATG-9 trafficking at the synapse, and ATG-9 subcellular localization, is associated with the promotion of local autophagosome formation. We hypothesize that the trafficking potentiates ATG-9 role as a lipid scramblase in the nucleation of the growth of the autophagosome isolation membrane^97–100^, perhaps via the activation of the scramblase activity or ATG-9-potentiated transport of lipids necessary for autophagosome biogenesis^103, 129^. Our observations that mutations in early autophagy protein EPG-9 similarly result in abnormal accumulation of ATG-9 in synaptic foci, which is enhanced by *cla-1(L)* mutants, underscore the relationship between ATG-9 exo-endocytosis at the synapse, active and periactive zone interaction, and autophagy.

**Figure 9.**
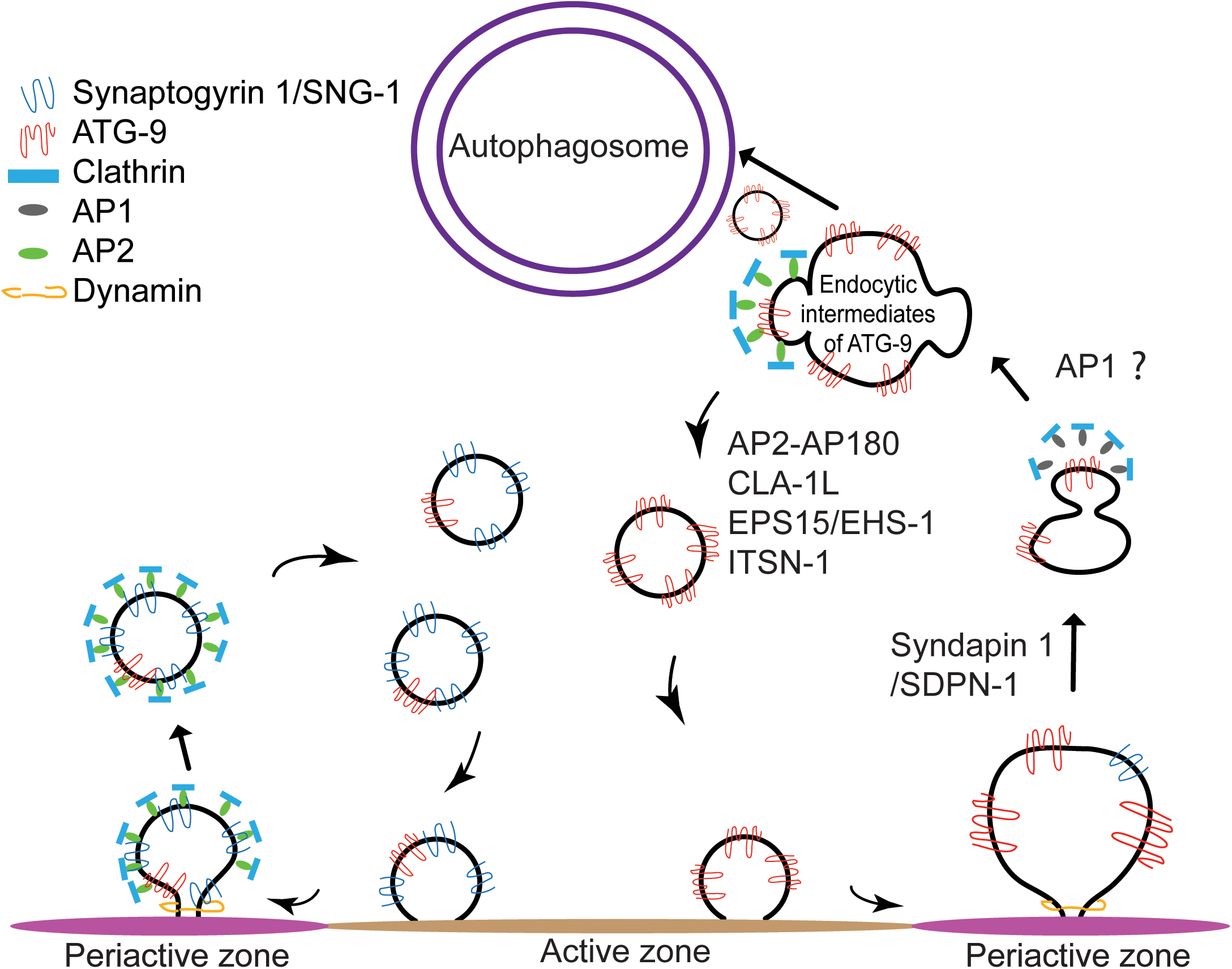
Cartoon diagram representing the genetic relationships between ATG-9 trafficking, the synaptic vesicle cycle and synaptic autophagy. At Zone 2 of AIY, both synaptic vesicles and ATG-9 vesicles undergo exo-endocytosis^21^. While most synaptic vesicles undergo clathrin-mediated endocytosis, our data are consistent with ATG-9 undergoing endosome-mediated endocytosis and displaying distinct phenotypes than those seen for synaptic vesicle proteins. We propose a model whereby ATG-9 is sorted by the adaptor complex AP1 to the endosome-like structures. CLA-1L, together with presynaptic endocytic proteins that reside in the periactive zone, such as EHS-1 and ITSN-1, as well as the adaptor complexes such as AP-2 and AP180, are necessary for sorting of ATG-9 from endocytic intermediates. Mutations in the active zone gene *cla-1L* result in abnormal accumulation of ATG-9 into endocytic intermediates, and defects in activity-dependent autophagosome formation.

ATG-9 trafficking at presynaptic sites via exo-endocytosis could link synaptic activity with local synaptic autophagy^21^, thereby modulating local autophagosome formation and degradation of synaptic proteins or organelles. Damaged synaptic proteins and vesicles associated with synaptic vesicle recycling can be degraded via autophagy^7, 110, 111, 130^, and ATG-9 trafficking has been associated with autophagosome biogenesis. Our observations are consistent with a model whereby the activity dependent trafficking of ATG-9 at the synapse via exo-endocytosis could track the activity state of the synapse, and instruct local autophagy to degrade damaged proteins or organelles in active synapses. Active zone proteins, like CLA-1L, which bridge the exocytic active zone regions with the endocytic periactive zone, could regulate ATG-9 trafficking to modulate this activity-dependent presynaptic autophagy.

## Materials and methods

### *C. elegans* strains and genetics

*C. elegans* Bristol strain worms were raised on NGM plates at 20°C using OP50 *Escherichia coli* as a food source^131^. Larva 4 (L4) stage hermaphrodites were examined. For a full list of strains used in the study, please see the KEY RESOURCES TABLE.

### Molecular biology and transgenic lines

Expression clones were made in the pSM vector^132^. The transgenic strains (0.5– 50 ng/μl) were generated using standard injection techniques and coinjected with markers Punc122::GFP (15–30 ng/μl), Punc122::dsRed (15–30 ng/μl) or Podr-1::rfp (15-30ng/μl). UNC-101, mouse AP1 mu1 and mouse AP2 mu isoform1 were PCR amplified from *C. elegans* and mouse cDNA (PolyATtract® mRNA Isolation Systems, Promega and ProtoScript® First Strand cDNA Synthesis Kit, NEB). Detailed sub-cloning information is available upon request.

### Forward genetic screen, SNP mapping and whole-genome sequencing (WGS)

*Cla-1(ola285)* was isolated from a visual forward genetic screen designed to identify mutants with abnormal localization of ATG-9::GFP in the interneuron AIY. *OlaIs34* animals, expressing ATG-9::GFP and mCherry::RAB-3 in AIY interneurons, were mutagenized with ethyl methanesulfonate (EMS) as described previously^131^ and their F2 progenies were visually screened under A Leica DM 5000 B compound microscope with an oil objective of HCX PL APO 63x/1.40-0.60 for abnormal ATG-9::GFP localization. Single-nucleotide polymorphism (SNP) mapping^45^ was used to map the lesion corresponding to the *ola285* allele to a 6.6-11.3 Mbp region on chromosome IV. Whole-genome sequencing (WGS) was performed at the Yale Center for Genome Analysis (YCGA) and analyzed on www.usegalaxy.org using “Cloudmap Unmapped Mutant workflow (w/ subtraction of other strains)” as described^46, 47^. The *ola285* allele was sequenced by using Sanger sequencing (Genewiz), and the genetic lesion confirmed as a single T-to-A nucleotide substitution in Exon 15 of *cla-1L* that results in a nonsense mutation I5753N. Complementation tests were performed by generating *ola285*/*cla-1(ok560)* trans-heterozygotes. The *ola285* allele failed to complement *cla-1(ok560)*.

### Cell autonomy of CLA-1L

A CRISPR protocol^133, 134^ was used to create *cla-1 (ola324)*, in which two loxP sites were inserted into two introns of *cla-1L*^40^. Cell-specific removal of CLA-1L in AIY was achieved with a plasmid driving the expression of Cre cDNA under the AIY-specific *mod-1* promoter fragment^135^.

### Cell autonomy and cell-specific rescues

The ATG-9 phenotype in *unc-101(m1);cla-1(ola285)* was suppressed by cell specifically expressing the *C. elegans* cDNA of *unc-101(m1)/AP1µ1*C, and the murine cDNA of *AP1µ1*. Cell-specific expression was achieved using AIY-specific ttx-3G promoter fragment^136^. Cell-specific expression of the murine cDNA of *AP2µ* in AIY did not suppress the ATG-9 phenotype in *unc-101(m1);cla-1(ola285)*.

### Confocal imaging

Hermaphrodite animals of Larval stage 4 (L4) were anesthetized using 10 mM levamisole (Sigma-Aldrich) or 50mM muscimol in M9 buffer, mounted on 2-5% agar pads and imaged by using a 60x CFI Plan Apochromat VC, NA 1.4, oil objective (Nikon) on an UltraView VoX spinning-disc confocal microscope (PerkinElmer). Images were processed with Volocity software. Maximum-intensity projections presented in the figures were generated using Fiji (NIH) for all the confocal images. Quantifications were performed on maximal projections of raw data.

### Quantification and statistical analyses

#### Quantifications of penetrance and expressivity

A Leica DM500B compound fluorescent microscope was used to visualize and screen the worms in the indicated genetic backgrounds. To quantify the penetrance, animals were scored as displaying either wild type (distributed throughout the Zone 2 synaptic region) or mutant (localized into subsynaptic foci) phenotypes for ATG-9 or SNG-1 at Zone 2 of AIY. Mutant phenotype was defined as one or more subsynaptic foci of ATG-9::GFP or SNG-1::GFP at Zone 2 of AIY. For each genotype, at least 40 animals were scored. The Y-axis of graphs is named as “% of animals with ATG-9 (or SNG-1) puncta at Zone 2 of AIY”. Statistics for penetrance quantification was determined using two-tailed Fisher’s exact test. Error bars represent 95% of confidence intervals.

To quantify the expressivity of the phenotypes, confocal micrographs of around 15 representative worms for each genotype were acquired using a spinning-disc confocal microscope (PerkinElmer). The Index of ATG-9 punctum at Zone 2 of AIY for each genotype was consistent across different confocal settings (data not shown), so ATG-9 subsynaptic foci in *cla-1(ola285)* were imaged with a different (lower exposure) confocal setting from the wild-type control, to avoid saturating the signal. Fluorescence values for each AIY Zone 2 were obtained after background subtraction by drawing a Freehand line using Fiji along the long axis of Zone 2. The fluorescence peak values and trough values were acquired via the Profile Plot function. Subsynaptic enrichment index was then calculated as (*Fpeak*-*Ftrough*)/*Ftrough*. The Y-axes of the graphs are named as “Index of ATG-9 (or SNG-1) punctum at Zone 2 of AIY”.

#### Mean intensity of SYD-2 at the presynaptic Zone 2 of AIY neurons

To measure the level of SYD-2 at presynaptic regions, we obtained the fluorescent value using Fiji as indicated above. All settings for the confocal microscope and camera were kept identical to compare the intensity of SYD-2 between the wild-type and *cla-1(ok560)* mutants (this calculation was different from that made for the ATG-9 index, where the (*Fpeak*-*Ftrough*)/*Ftrough* was calculated within the same synapse). The same ROI was drawn for all micrographs analyzed. The mean fluorescent value of SYD-2 was measured by Fiji.

#### Colocalization analysis

The Coloc2 plugin of Fiji was used to measure the Pearson correlation coefficient for colocalization between ATG-9::GFP and CHC-1::BFP, or ATG-9::GFP and SNG-1::BFP, both in *cla-1(ola285)* mutants. ROI was drawn to include the entire Zone 2 in all Z-stacks.

#### Activity-dependent autophagy

To measure the synaptic autophagosomes in AIY, *olaIs35* animals expressing eGFP::LGG-1 and cytoplasmic mCherry in AIY interneurons in the wild-type, *cla-1(ola285)* and *cla-1(ok560)* mutant backgrounds were raised in 20°C and then shifted to 25°C for 4 hours as described^21^ to alter the activity state of the AIY interneuron^95^. LGG-1 puncta in the neurite of AIY was scored under a Leica DM 1048 5000B compound microscope as described^15^.

#### Statistical Analyses

Statistical analyses were conducted with Prism 7 software and reported in the figure legends. Briefly, two-tailed Fisher’s exact test was used to determine statistical significance of categorical data such as the penetrance of ATG-9 phenotype. For continuous data, such as Index of ATG-9 punctum, two-tailed unpaired Student’s t-test or ordinary one-way ANOVA with Tukey’s post-hoc analysis was used. For LGG-1 scoring, since the number of LGG-1 puncta for each condition did not pass D’Agostino & Pearson normality test, the nonparametric analysis Kruskal-Wallis with Dunn’s multiple comparisons test, was used. The error bars represent either 95% confidence interval or standard deviation (SD), as indicated in the figure legends, along with the p values.

### Protein sequence alignment of the AP adaptor complex *µ* subunit

The sequence of *C. elegans* unc-101/AP-1µ1 (K11D2.3) was acquired from WormBase (https://wormbase.org). The sequences of murine AP-1µ1 (NP_031482.1) and murine AP-2µ (NP_001289899.1) were acquired from NCBI (https://www.ncbi.nlm.nih.gov/protein/?term=). Sequence alignment of worm UNC-101/AP1µ1, mouse AP1µ1 and mouse AP2µ was generated using Tcoffee in Jalview 2.10.5^137^.

## KEY RESOURCES TABLE

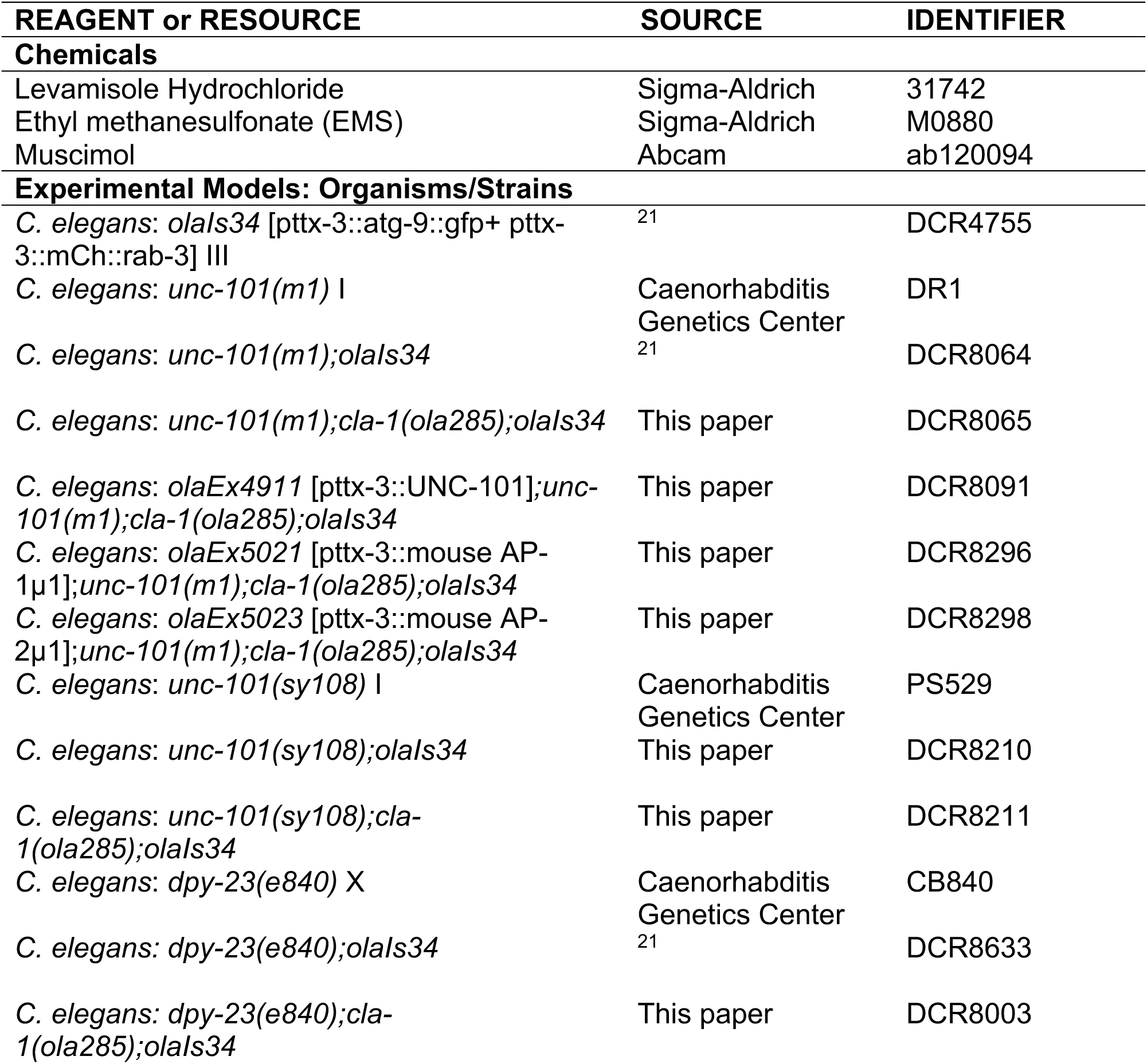
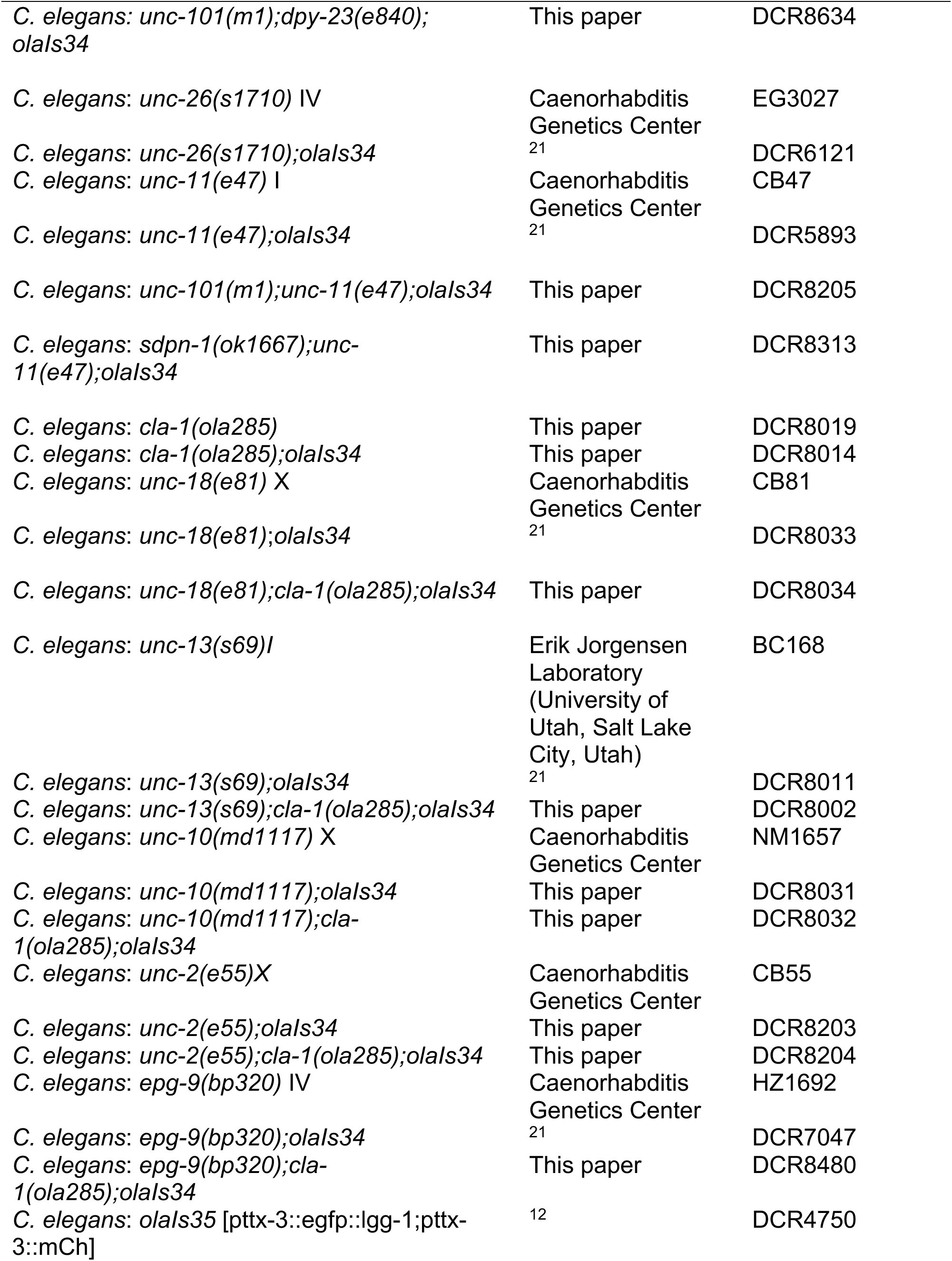
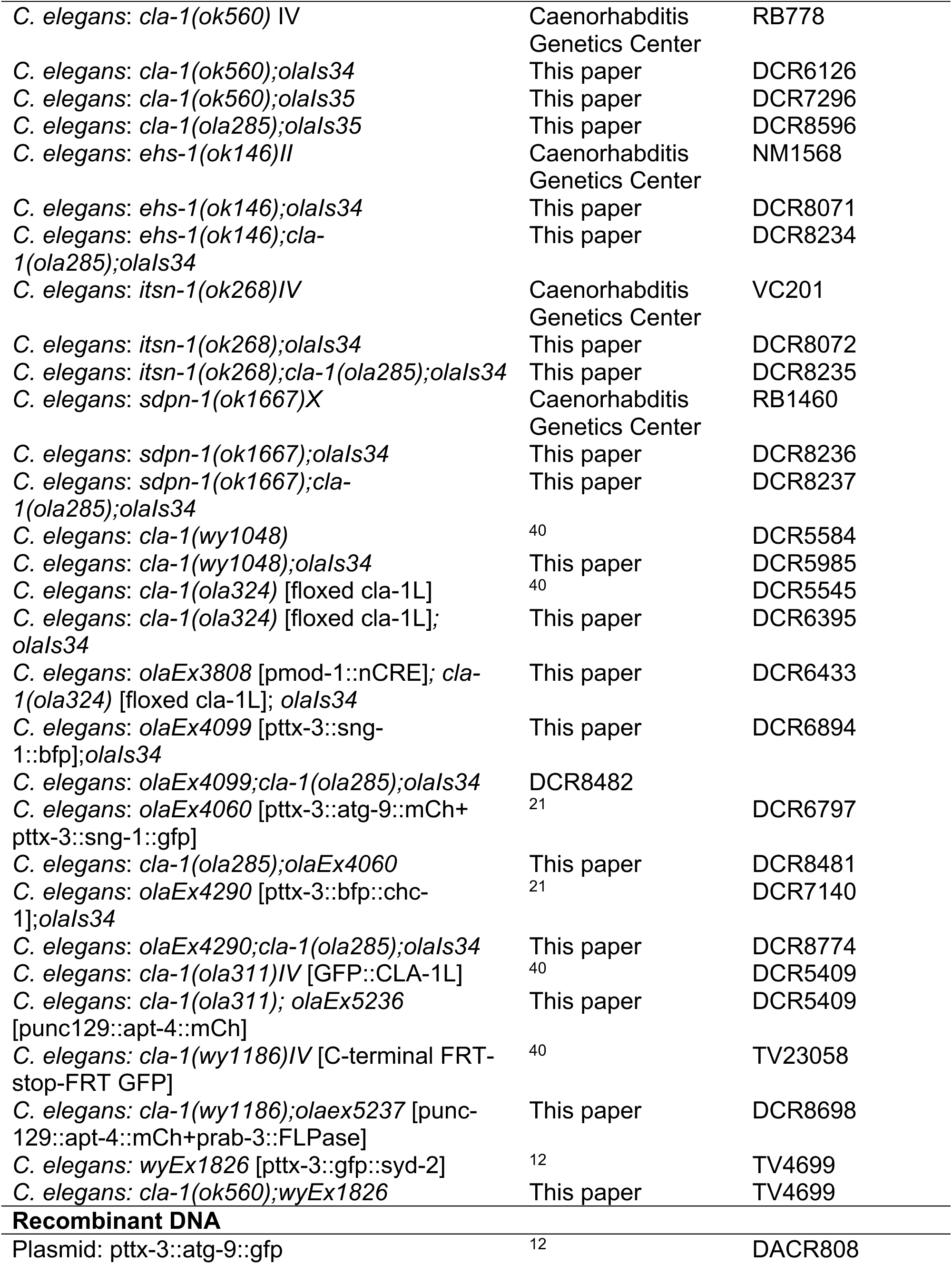
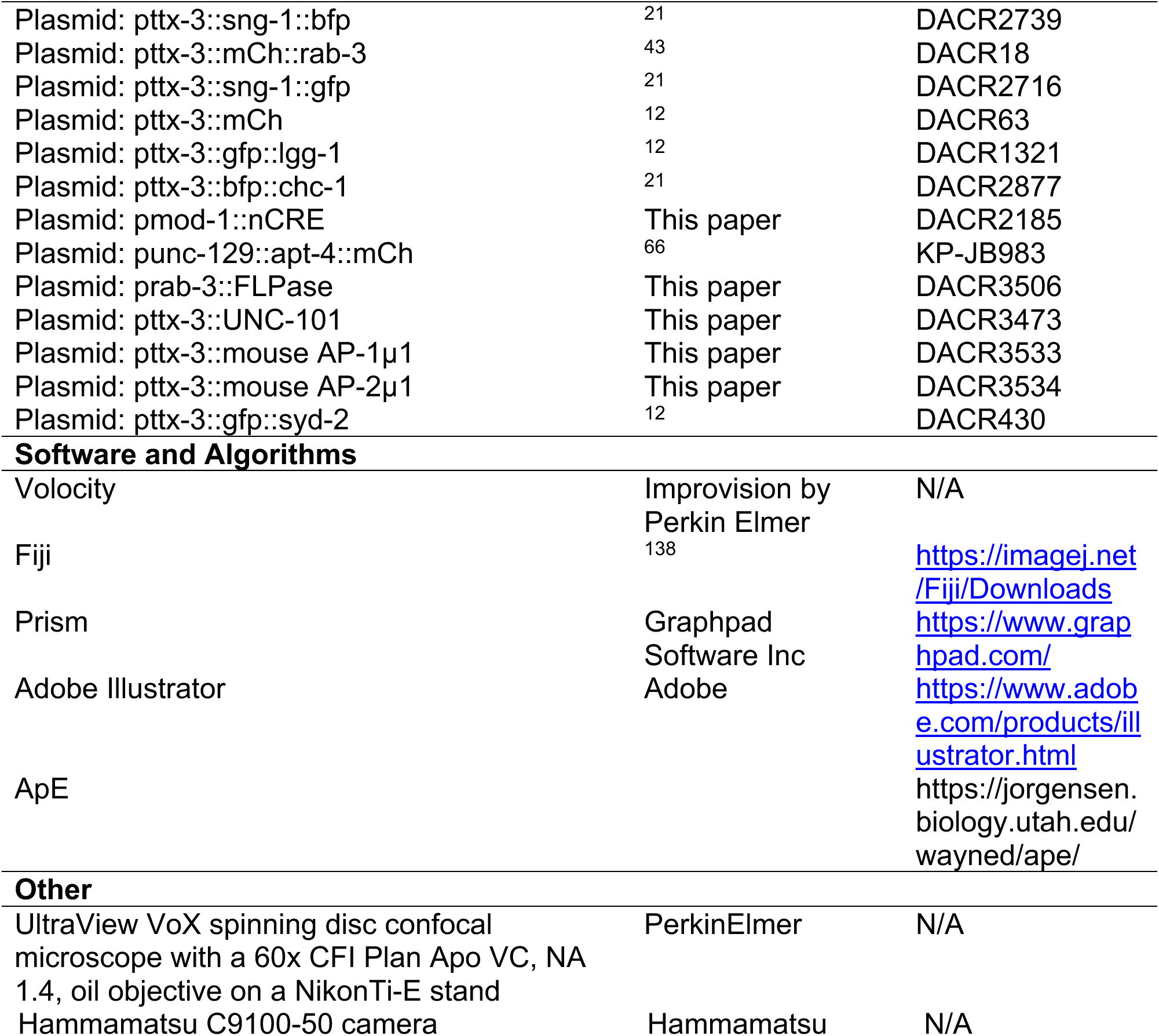

## Acknowledgments

We thank Jihong Bai (Basic Sciences Division, Fred Hutch) and Kang Shen (Department of Biology, Stanford University) for providing strains and constructs. We thank Lin Shao (Department of Neuroscience, Yale University) for assistance with image quantification and statistics. We thank Josh Hawk for providing mouse cDNA. We thank current and past members of the Colón-Ramos lab for help, advice and insightful comments on the project. We also thank Andrea Stavoe, Mia Dawn, Peri Kurshan, Janet Richmond and Pietro De Camilli for comments on the project. We thank the Caenorhabditis Genetics Center (funded by NIH Office of Research Infrastructure Programs P40 OD010440) for *C. elegans* strains. S.Y. was supported by China Scholarship Council-Yale World Scholars Program. Research in the D.A.C.-R. lab was supported by NIH R01NS076558, DP1NS111778 and by an HHMI Scholar Award.

## Contributions

Z. X., S.Y., and D.C.R. conceived the study and designed the experiments. Z.X., S.Y., and S.E.H. conducted the experiments. Z.X. performed image quantification and biostatistics analysis. Z.X. and D.C.R. prepared the manuscript with contribution from S.Y., S.E.H., B.C., and L.M.

---

**Supplemental figure 1.**
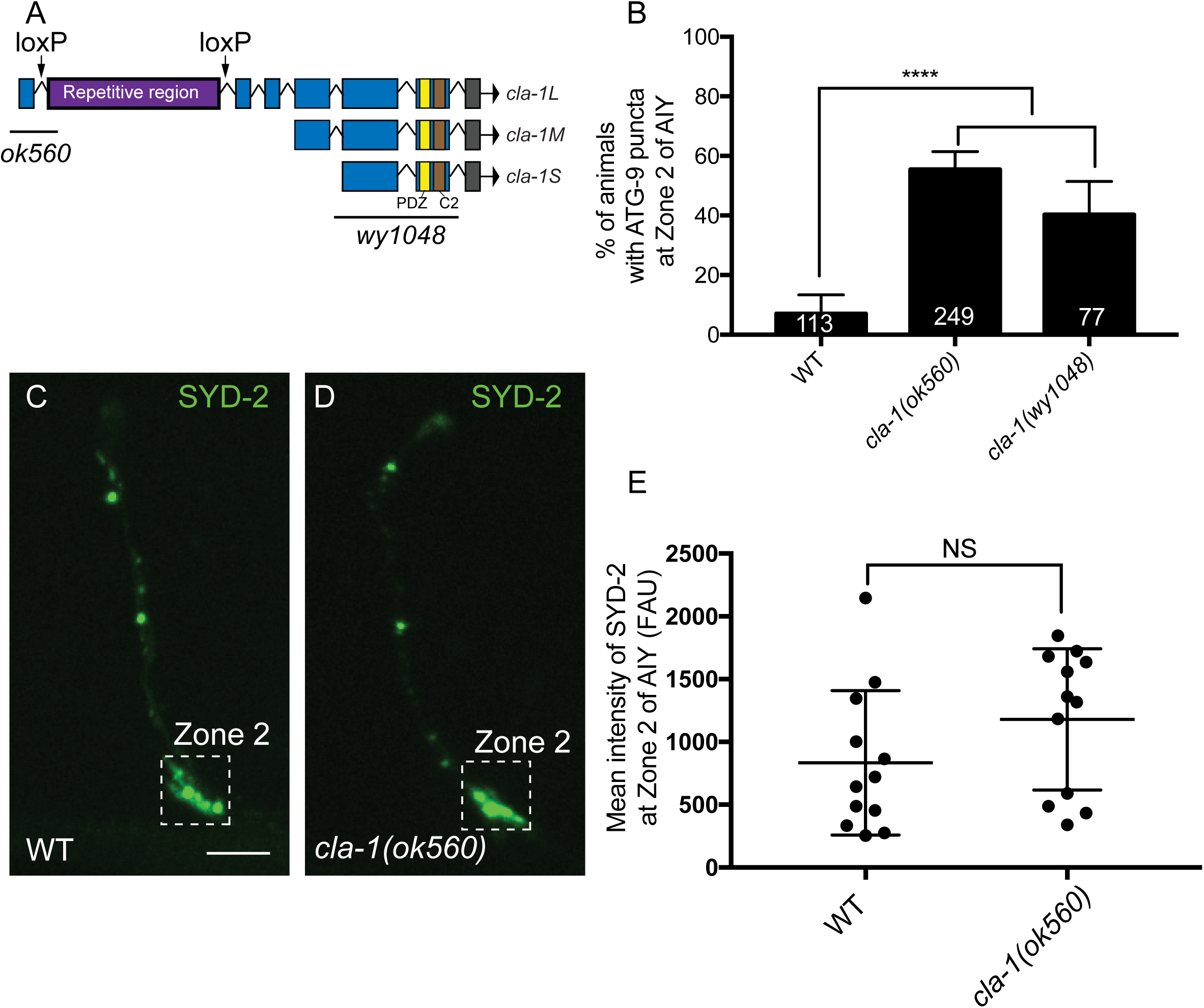
Examination of the active zone protein SYD-2 in *cla-1(L)* mutants, and of ATG-9 distribution in *cla-1(wy1048)* null allele. (A) Schematic of *cla-1* gene, with different protein isoforms. The *ok560* allele specifically affects the long protein isoform, while *wy1048* allele affects all CLA-1 protein isoforms. (B) Quantification of the percentage of animals displaying ATG-9 subsynaptic foci at AIY Zone 2 in the indicated genotypes. Error bars represent 95% confidence interval. ****p<0.0001 by two-tailed Fisher’s exact test. The number on the bars indicates the number of animals scored. (C-D) Distribution of SYD-2::GFP at the synaptic Zone 2 and Zone 3 regions of AIY in wild type (C) and *cla-1(ok560)* (D) animals. The dashed boxes highlight the presynaptic Zone 2 of AIY examined in this study. (E) Mean intensity of SYD-2 at AIY Zone 2 in wild type (WT) and *cla-1(ok560)* mutants. Error bars show standard deviation (SD). “NS” (not significant) by two-tailed unpaired Student’s t-test. Each dot in the scatter plot represents a single animal. Scale bar (in C for C-D), 5 μm.

**Supplemental figure 2.**
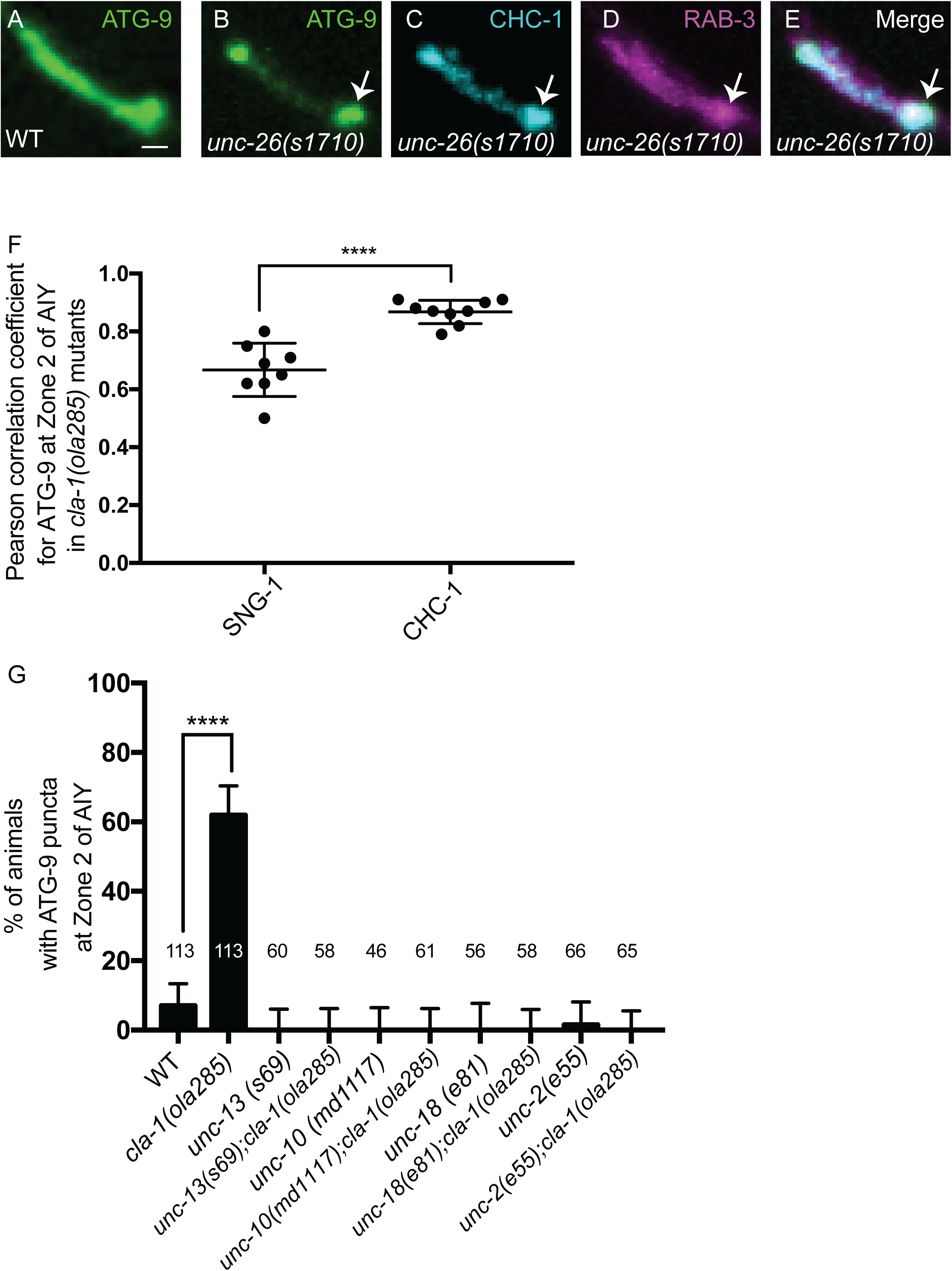
ATG-9 accumulates at clathrin-rich foci in *cla-1(ola285)* mutants, which is suppressed by mutants for synaptic vesicle exocytosis. (A) Distribution of ATG-9::GFP at Zone 2 of AIY in wildtype (WT) animals (A). (B-E) Distribution of ATG-9::GFP (B), BFP::CHC-1 (pseudo-colored cyan) (C) and mCherry::RAB-3 (pseudo-colored magenta) (D) at Zone 2 of AIY (merge in E) in *26(s1710)/synaptojanin 1* mutant animals (see also ^21^). ATG-9 subsynaptic foci (indicated by arrows in B-E) are enriched with CHC-1 in *unc-26(s1710)/synaptojanin 1* mutants. (F) Pearson correlation coefficient for colocalization between ATG-9::GFP and SNG-1::BFP ,or between ATG-9::GFP and BFP::CHC-1, both in *cla-1(ola285)* mutants. Error bars show standard deviation (SD). ****p<0.0001 by two-tailed unpaired Student’s t-test. Each dot in the scatter plot represents a single animal. (G) Quantification of the percentage of animals displaying ATG-9 subsynaptic foci at AIY Zone 2 in the indicated genotypes. Error bars represent 95% confidence interval. ****p<0.0001 by two-tailed Fisher’s exact test. The number on the bars indicates the number of animals scored. Scale bar (in A for A-E), 1 μm.

**Supplemental figure 3.**
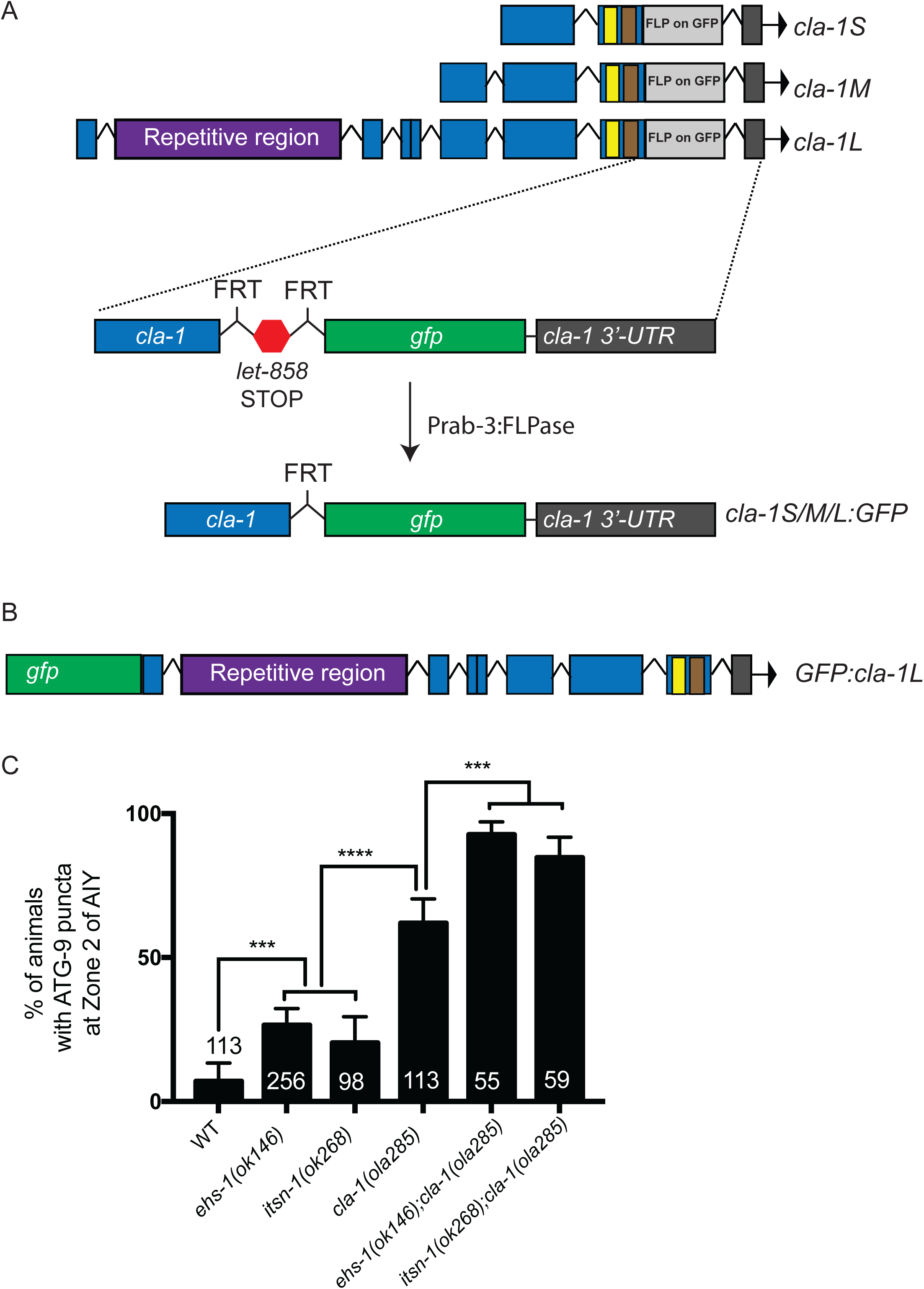
CLA-1L genetically interacts with the endocytic proteins at the periactive zone to regulate ATG-9 trafficking. (A) Schematics of the strategy for endogenously tagging CLA-1 at C-terminus via CRISPR. FLP-on-GFP (let-858 3’UTR flanked by FRT sites followed by GFP) was inserted at the common C-terminus of CLA-1 (shared by CLA-1L, CLA-1M and CLA-1S protein isoforms, see ^40^). FLPase driven by the Prab-3 promoter is expressed pan-neuronally to induce expression of CLA-1::GFP in an endogenous manner (see Figure 4A). (B) Schematics of endogenously tagging CLA-1L at N-terminus via CRISPR ^40^. GFP was inserted at the unique N-terminus of CLA-1L (see Figure 4H). (C) Quantification of the percentage of animals displaying ATG-9 subsynaptic foci at AIY Zone 2 in the indicated genotypes. Error bars represent 95% confidence interval. ***p<0.001, ****p<0.0001 by two-tailed Fisher’s exact test. The number on the bars indicates the number of animals scored.

**Supplemental figure 4.**
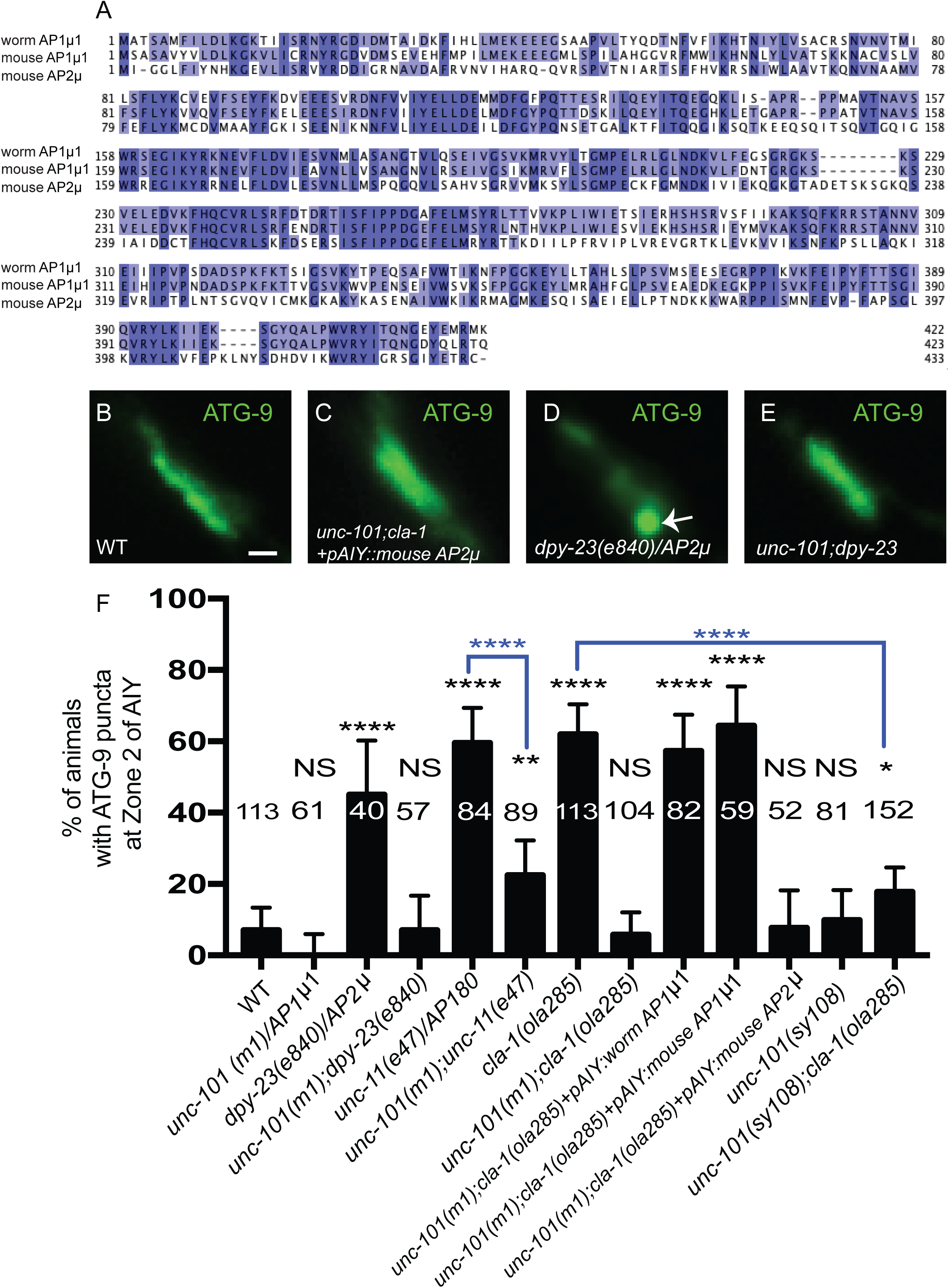
The AP-1 and AP-2 adaptor complexes mediate presynaptic trafficking of ATG-9. (A) Sequence alignment of *C. elegans* UNC-101/AP1µ1, mouse AP1µ1 and mouse AP2µ, generated using Tcoffee in Jalview^137^. Although both AP-1 µ1 and AP-2µ share similarity, *C. elegans* UNC-101/AP1µ1 is more similar to the mouse AP1µ1 (Query Cover: 100%; Percentage Identity: 74%) than the mouse AP2µ (Query Cover: 98%; Percentage Identity: 41%). (B-E) Distribution of ATG-9::GFP at Zone 2 of AIY in wild type (WT) (B), *unc-101(m1);cla-1(ola285)* mutants with mouse AP2µ cDNA cell-specifically expressed in AIY (C), *dpy-23(e840)/AP2µ* (D) and *unc-101(m1);dpy-23(e840)* (E) mutant animals. ATG-9 subsynaptic foci are indicated by the arrow (in D). (F) Quantification of the percentage of animals displaying ATG-9 subsynaptic foci at AIY Zone 2 in the indicated genotypes. Error bars represent 95% confidence interval. “NS” (not significant), *p<0.05, **p<0.01, ****p<0.0001 by two-tailed Fisher’s exact test. The number on the bars indicates the number of animals scored. Black asterisks indicate comparison between each group with the wild-type control. Blue asterisks indicate comparison between two specific groups (highlighted with brackets). Scale bar (in B for B-E), 1 μm.

**Supplemental figure 5.**
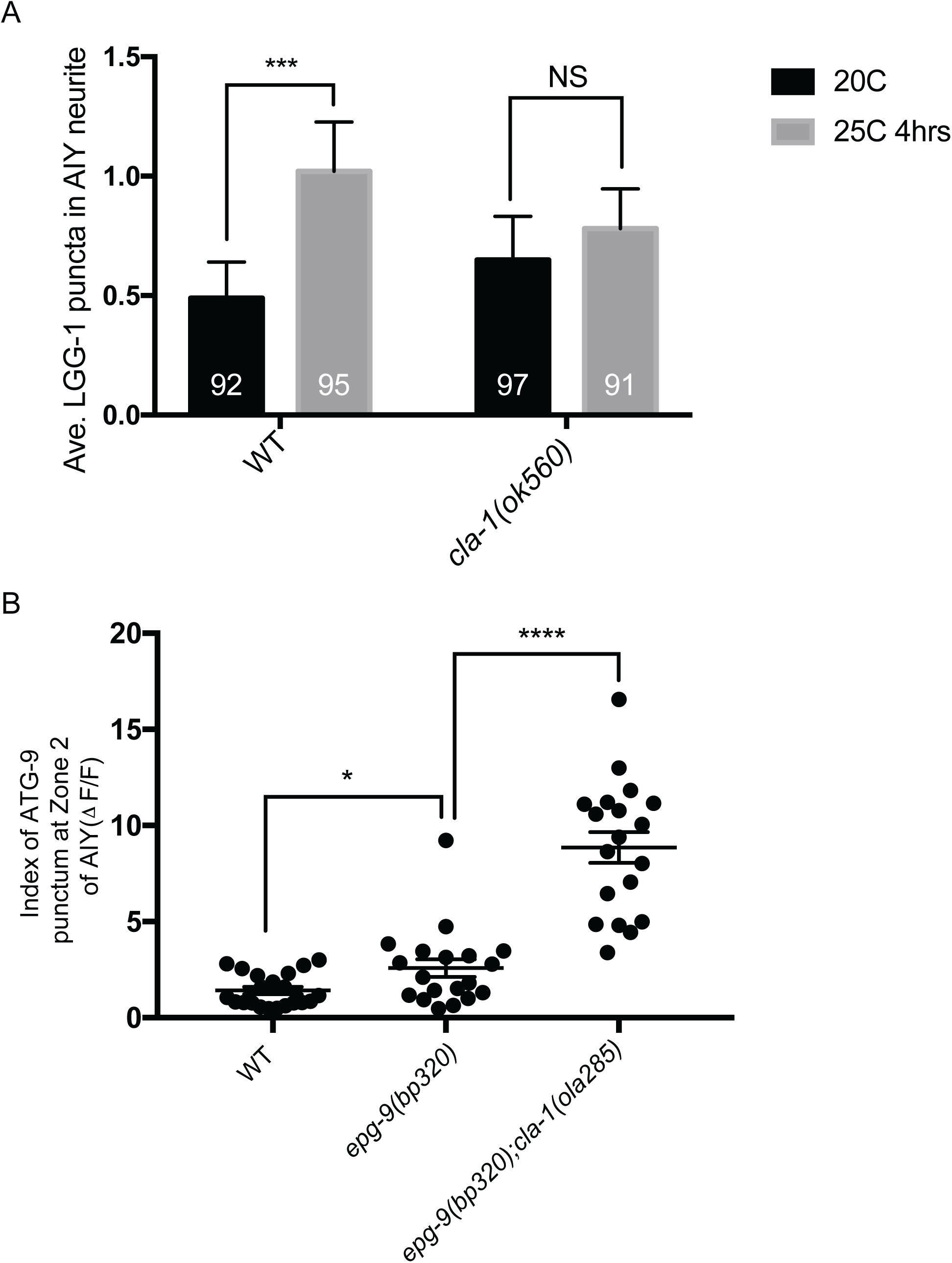
Disrupted ATG-9 trafficking in *cla-1(L)* mutants is associated with a deficit in activity-induced autophagosome formation. (A) Quantification of the average number of LGG-1 puncta in the AIY neurites at 20°C and at 25°C for 4 hours in wild type (WT) and *cla-1(ok560)* mutants. Error bars represent 95% confidence interval. “NS” (not significant), ***<0.001 by Kruskal-Wallis test with Dunn’s multiple comparisons test. The number on the bars indicates the number of animals scored. (B) Quantification of the index of ATG-9 punctum (ΔF/F) at Zone 2 of AIY in the indicated genotypes. Error bars show standard deviation (SD). *p<0.05, ****p<0.0001 by ordinary one-way ANOVA with Tukey’s multiple comparison test. Each dot in the scatter plot represents a single animal.

